# S-SELeCT: A Human-Evolved Serine Integrase System for Efficient Large-Cargo Genome Integration

**DOI:** 10.64898/2026.01.30.702954

**Authors:** Alfonso P. Farruggio, Lin Jiang, Karen Duong, Cynthia Nguyen, Razan Kaddoura, Ruby Tsai

## Abstract

As a consequence of their sizes, many loss-of-function genetic mutations fall within large genes. A major gene-therapy tool that could be used to solve large swaths of the genetic diseases that result from these inherited mutations is large-fragment knock-in. I.e. instead of attempting to create separate treatments for each and every location that these mutations occur in, large groups of patients could be aided via a single safe-harbor integration of the full-length coding sequence. Towards this goal, we have created a set of early stage gene-editing enzymes that can help mediate large cargo integration at a safe harbor locus in human cells. When expressed in stable lines, our S-SELeCT (<u>S</u>ite-<u>S</u>pecific <u>L</u>arg<u>e C</u>argo <u>T</u>argeting) integrase fusions can facilitate integration of a 10 kb plasmid at frequencies up to 32%, and when delivered transiently via plasmid transfection, we were able to achieve up to 13% knock-in. These are the first serine integrase enzymes that have been evolved fully in human cells, and the first to recognize an endogenous symmetric non-pseudosite – the first true human serine integrase attachment site.

**GRAPHICAL ABSTRACT:** 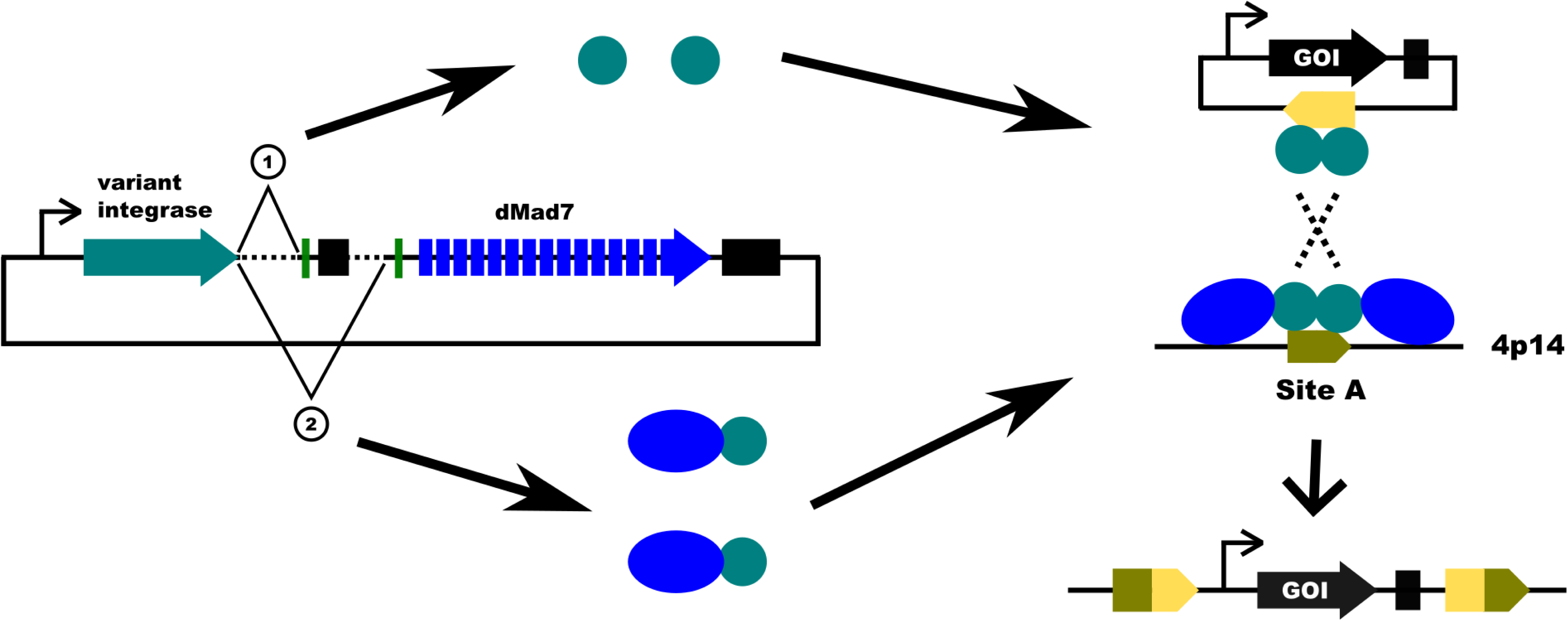

## INTRODUCTION

Since the initial reports of eukaryotic gene editing in 2012-2013, the CRISPR/Cas9 system has completely changed the ease and speed at which cell line and animal models are developed (1), and CRISPR-based therapeutics have now been part of hundreds of clinical trials. However, while CRISPR performs exceptionally well at cutting DNA and knocking out genes, Cas9-mediated transgene knock-in often suffers from low efficiencies, due to a reliance upon the endogenous host cell’s repair machinery for the introduction of exogenous DNA. In addition, these DNA breaks can lead to undesired outcomes like off-target indels, large deletions, translocations and inversions (2–5). Therefore, there is a need for technologies that can drive site-specific insertion of large transgenes for therapeutic gene editing.

Serine integrases are excellent candidates for this because they facilitate both the cleavage and re-ligation reactions that are needed for transgene knock-in (6), thereby eliminating reliance upon host DNA-repair machinery. These enzymes mediate integration via recombination of two distinct 40-50 bp sequences called the attP and attB sites, which are named after on the locations that they naturally occur in: the phage and bacterial attachment sites, respectively (6). In the absence of the Recombination Directionality Factor (RDF) protein, this reaction is unidirectional, i.e. dimers bound to the resultant attL and attR sites (left and right attachment sites) do not form tetramers, thus the reverse reaction cannot proceed (6, 7). Att-sites have dyad symmetry, i.e. they are semi-identical inverted repeats that bind to homodimers of the integrase protein (6, 7).

While serine integrases have evolved to function in prokaryotes, a variety have been observed to function well in mammalian cells (8, 9). Some of these enzymes can mediate integration into endogenous mammalian sequences called “pseudosites”, however most require that their wildtype attP or attB sequence be pre-introduced into the genome, especially for optimal integration efficiency (9, 10). Unlike recombination between their natural sequences, pseudosite integration often involves host-cell DNA-repair activities, which can result in undesired outcomes like small insertions and deletions (indels) near the insert junctions, but also larger deletions, translocations and inversions (10, 11). Pseudosites are often not symmetrical, and usually bear only low-levels of identity to wildtype attP and attB sites (10–12).

To advance large-insert gene-therapy, we have derived S-SELeCT integrase enzymes, which are variants of phiC31 integrase (C31-int) created via directed evolution. With assistance from fused dMad7, a deactivated Cas nuclease (13, 14), they are capable of mediating plasmid knock-in to a novel human safe-harbor locus on chromosome 4 (4p14) that we refer to as “site A”. While many C31-int attP pseudosites are present in the human genome, integration frequencies at these sites is far lower than that for introduced wildtype attP (9, 15, 16). Thus, to more closely mirror the wildtype reaction, which facilitates high-efficiency no-damage recombination (15, 16), we have opted to evolve C31-int variants that target a symmetric non-pseudosite. I.e. site A is not a known pseudosite, and it has dyad symmetry that falls within the range of standard wildtype serine integrase attP sequences (Fig. 1).

**Figure 1:**
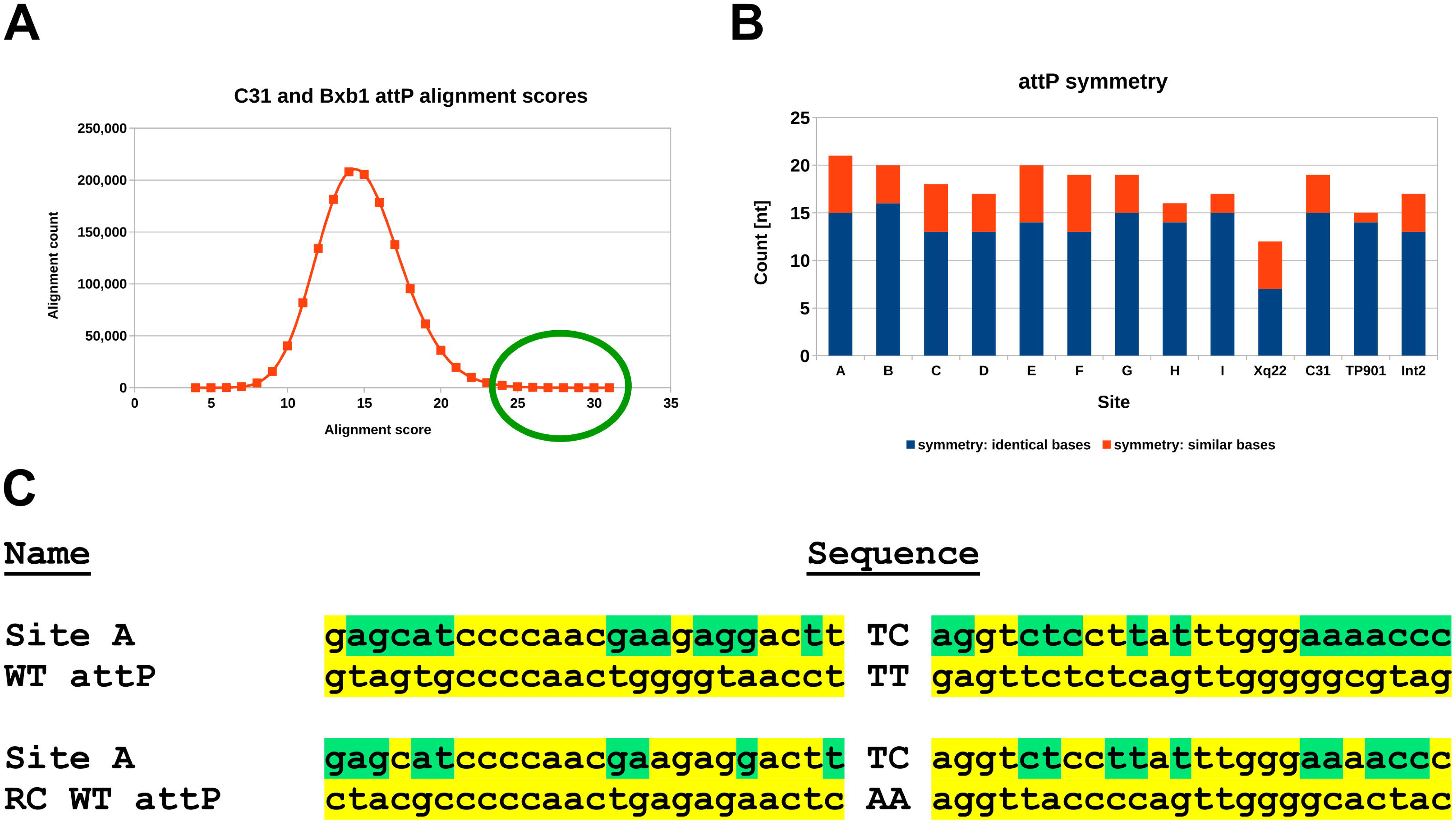
Site candidate search: Alignment scores and symmetry (**A**) attP-like sites’ alignment against C31-int attP. Approximately 75 sites, exhibiting the highest alignment scores, were subjected to further analysis (green circle). (**B**) Site symmetry counts (left vs right half-site alignment results) for candidate-sites A-i, pseudosite Xq22 and three wildtype serine integrases. Blue shows identical bases, and red shows similar bases (i.e., A is similar to G, and C is similar to T). (**C**) Sequence alignments of Site A and both orientations of WT C31-int attP (RC: reverse complement). In the Site A sequence, yellow bases match C31-int attP, while green bases do not.

To our knowledge, while there have been several previous attempts to alter the specificity of C31-int, they have all only yielded variants that performed well in bacteria. I.e. when the top mutants were tested in mammalian cells, their activity was majorly reduced (e.g. Sclimenti et al., Farruggio and Calos, and personal communications with Calos) (17, 18). Thus, to avoid these problems, we have performed all S-SELeCT variant screens in HEK293 cells. Screening in mammalian cells is preferable to bacteria or other organisms since activity in human cells is our primary interest, and it is not unusual for wildtype or variant prokaryotic DNA-modification enzymes to perform significantly worse in mammalian cells (8, 18, 19). Despite having the same sequence, a protein expressed in bacteria and mammalian cells can have significantly distinct activity because of different host-cell organelles (protein localization), post-translational modification systems (phosphorylation, glycosylation, etc.), and even protein-folding chaperones (20–22). Utilization of a mammalian-cell-based evolution system ensures that we are evolving toward a final protein that will function in mammalian cells.

## MATERIAL AND METHODS

### Deactivated Mad7

Since no precedence for the use of deactivated Mad7 (dMad7) in mammalian cells existed at the time we performed this work, we evaluated the three mutations described by Price et al.: D877A, E962A, and D1213A (14). The Mad7 sequence from Wierson et al. was modified to create four dMad7 candidates: d1a (D877A), d1b (E962A), d2 (D877A, E962A) and d3 (D877A, E962A, D1213A) (13). A custom EGxxGFP reporter assay (e.g. see Mashiko et al.) (23) was used to compare activity of the candidates to active Mad7 (data not shown). d2 and d3 triggered the least homologous-recombination, so they were next tested in triplicate, where d2 was observed to be optimal (i.e. least activation of GFP; Fig. S1). We thus adopted d2-Mad7 for use in all integrase-dMad7 fusions.

### Alternative-splicing WT C31-int-GFP/mCherry plasmid cloning

The plasmids used to test alternative-splicing introns were assembled using GGA of synthesized DNA segments and PCR products. They consist of two divergent promoterless expression cassettes centered on a Bxb1 attB site. The first incomplete cassette contains the coding sequence for a blasticidin-resistance protein, and it is terminated by a BGH 3’UTR/polyA. The second partial-cassettes contain the following elements: intronized WT C31-int, splice donor, splice-1 acceptor, P2A peptide, EGFP4m, splice-2 acceptor, P2A peptide, mCherry2, WPRE, SV40 3’UTR/polyA. In addition, we cloned a version of all plasmids with a BGH 3’UTR/polyA between EGFP4m and the splice-2 acceptor (“with internal polyA”, (Fig. S2). The alternative splice donors and acceptors are all derived from chicken TNNT intron 4 as described by Aebischer-Gumy et al. (24). In addition, two UMS terminators are present in the plasmid, one on each flank of the divergent partial-cassette segment. We have deposited sequences of the I4(7Y)-I4sh constructs with (pLK-int-GmC-4) and without (pLK-int-GmC-4-alt) an internal polyA at Zenodo.

### Transient fused-integrase expression plasmid assembly

Precursors of plasmids used to express integrase-dMad7 fusions were made using GGA of synthesized DNA and PCR products. They consist of three cassettes that express: (i) g6 gRNA, (ii) g10 gRNA, and (iii) an intronized integrase-dMad7 fusion with or without alternative-splicing. The gRNA-expression cassettes are driven by the human U6 promoter, and integrase-fusions are expressed using a CAG or mPGK promoter. In some plasmids, a WPRE and rabbit beta-globin 3’UTR/polyA are used, while in others just a BGH 3’UTR/polyA is present. All utilized introns and alternative splicing sequences match what were used in the respective stable-expression vectors. We have deposited the sequences of precursor plasmids (pLK-CAG-357-NL1-5C, pLK-CAG-357-NL2-5C, pLK-mPGK-357-NL1-5C, pLK-mPGK-357-NL2-5C, pLK-CAG-357-NL1a4-5C, pLK-CAG-357-NL2a4-5C, pLK-mPGK-357-NL1a4-5C-W, pLK-mPGK-357-NL2a4-5C-W), and two alternative-splicing vectors (pLK-CAG-V12-NL1a4-5C, pLK-mPGK-V7-NL2a4-5C-W) at Zenodo.

Vectors used to compare transient expression of WT C31-int fusions to dCas9a (25) and dMad7 (Fig. S3) were created via GGA of synthesized DNA segments and PCR products. They consist of a single CMV-driven cassette with a polycistron, 5’ HIV1 TAR (26, 27), 3’ WPRE, and that is terminated by a BGH 3’UTR/polyA. The polycistrons encode HIV1-Tat (26, 27), a C31-int-dCas protein, and mCherry 2. The three proteins are separated via 2A skipping peptides (P2A and T2A). C31-int, dCas9a and mCherry2 are heavily intronized in all constructs, whereas dMad7 was only cloned this way in one construct (Fig. S3). We have deposited the sequences of these constructs (pLKCTT-i23-L8-miC-W, pLKC-L8-dM7-TTF, pLKC-L8-i20-dM7-TTF) at Zenodo.

### Mutagenesis of C31-int to create variant libraries

Integrase-variant libraries were generated using a variety of methods, including regional error-prone PCR (e.g. Yang et al.) (28), multi-codon saturation, and GGA-shuffling (29) of segments containing mutations from enriched pools. For codon saturation involving oligos, we used NNK degeneracy (30). In oligo pools, in addition to NNK, we sometimes synthesized 20 separate oligos (one for each possible amino acid) to perform saturation. Insertion and deletion saturation involving 1-4 codons was also performed using oligo pools. For insertion saturation, a combination of NNK and separate-oligos were used to vary amino acids that were introduced, but also at the flanking positions. When performing deletions, the flanking positions were saturated using NNK and separate-oligos in the pool. All synthetic DNA used for mutagenesis was purchased from IDT (oligos, Ultramers and oPools), and GGA was always used to fuse mutant segment pools with plasmid backbones.

Mutation summaries for V7, V12, V29, V30, V32, V36, V37, and V40, which were top-performers in our most difficult assays, have been provided in supplementary tables S1-S8. Total mutation counts for these variants are listed in Table S9.

### Creation of the HEK293 H11 landing-pad line

Our landing-pad cell line was created by knocking-in a Bxb1 integrase landing-pad (LP) at the H11 locus in a HEK293 line that stably expresses a doxycycline-activated transcription factor (31). To introduce the LP, we used the same TALEN-stimulated approach described in our DICE publication (15). A plasmid mix that consisted of 833 ng MR015-H11-L2-TALEN (Addgene 51554), 833 ng MR015-H11-R2-TALEN (Addgene 51555), and 1665 ng of the LP plasmid was complexed with 1 uL Xfect, then 2 ug of it was delivered into a 24 ww. After 72 hours, we expanded the cells 1:20 and grew them under 200 ug/mL hygromycin selection for 2 weeks. Single cells from this pool were sorted via FACS, and outgrowths were genotyped for the desired knock-in. We have deposited the Tet promoter-attP-EF1a promoter LP segment sequence depicted in Fig. S4A at Zenodo (H11_LP_core).

### Establishment of solo or fused-integrase variant HEK293 libraries

To create pools of HEK293 cells that express a single solo or fused integrase variant from the H11 locus, we co-transfected a Bxb1-integrase expression vector with a stable-expression plasmid library into our LP cell line. Our primary Bxb1 integrase plasmid contains a codon-optimized sequence with a C-terminus X. laevis nucleoplasmin NLS; the vector (pCMV-opt-Bxb1-XeNLS) sequence has been deposited at Zenodo. For transfection into a well of a 6-well plate (6ww), the DNA mix contained 2.75 ug Bxb1 integrase vector, 8.25 ug variant-integrase plasmid pool, and was brought to a total volume of 179.8 uL with Xfect Buffer. After complexing with 3.3 uL Xfect, 10 ug (166.7 uL) was delivered into the 6 ww. In parallel, to estimate library size or coverage, a promoterless attB-mCherry donor was co-transfected with the Bxb1 integrase into separate wells (seeded the same way). Three days after transfection, library size or coverage was estimated by measuring the percent of mCherry-positive cells above background (integration efficiency) in our attB-mCherry control (e.g. Fig. S4B). I.e. library diversity = integration efficiency x total cell count. If the library size and coverage (= diversity / size) was adequate, the cells were expanded 1:20 and maintained for two weeks in medium supplemented with 20 ug/mL blasticidin. The cells were fed 1-3 times a week, and split as needed to maintain at least 3X library coverage.

## RESULTS

### Selection of site A

To search for attP-like sites for our evolution work, we employed the following criteria: (1) putative sites should have high sequence homology to wild-type C31-int attP, (2) they should possess similar degrees of dyad symmetry that are commonly observed in wild-type attP sites, and (3) they must be located in an intergenic region (i.e. we do not want to disrupt a gene). Having a high sequence-similarity to C31-int attP helps to decrease the number of directed-evolution cycles needed, and increases our likelihood of success. Symmetry is important because homodimers (identical integrase molecules bound via their catalytic domains) bind to att-sites. As shown in Fig. 1, attP sites from C31-int, TP901, and Int2 have similar degrees of symmetry, much higher than that of a well-studied pseudo-site, Xq22.1, which has previously performed poorly in directed evolution attempts.

A custom program was written to screen the entire human genome for regions exhibiting the dyad symmetry that is found in wildtype (WT) serine integrase attP sites. Over 1 million sites were returned, and identified genomic sites were plotted as a function of alignment score to C31-int and Bxb1 integrase attP sequences (Fig. 1A). Approximately 75 sites, exhibiting the highest alignment scores (green circle in Fig. 1A), were further analyzed to identify sequences which met the intergenic location requirement outlined above. In addition, the following criteria were applied: (a) sites should have a GC content between 20-60%, which is the range typically observed for wild-type serine integrases, and (b) sites should be unique (e.g. not present in repeat elements).

We focused on C31-int candidates because they were the most promising – putative human attP sites, A through i, are listed in Fig. S5, along with their locus information (chromosome arm and band), the neighboring 5’ and 3’ genes, and their distance from the site. Further, the symmetry scores for these sites (Fig. 1B) are all similar to three representative wild-type attP sites bound by the phiC31, TP901, and Int2 serine integrases. The sites were prioritized based on proxies for chromatin accessibility: (1) evidence for active expression from neighboring genes, and (2) proximity to DNAseI hypersensitive sites (HSS). Chromatin accessibility was further validated for sites A, C, D and E by confirming that Cas9 could mediate dsDNA integration at the loci in HEK293 cells (Table S10 and data not shown).

We chose Site A as the directed evolution target because of its homology, symmetry, and location. Compared with WT phiC31 attP, 22 and 29 of the 48 nucleotides are identical in the forward and reverse-complement alignments, respectively (i.e. di-nucleotide core not considered; Fig. 1C). Considering that it would likely be difficult to directly identify a C31-int variant that recognizes Site A in a single round of screening, we instead followed the example of Sarkar et al. (32). I.e., a series of intermediate attP sequences were generated in order to allow us to gradually evolve C31-int from WT to Site A attP and attB recognition.

### Directed evolution using the inversion assay screen

Site A was divided into 5 segments, A1-A5 (Fig. 2), each of which differs from C31 attP by 3-4 bases in each half site (6-8 bases total) (Fig. S6), and are grouped based on predicted shared proximity to key amino acids in bound C31-int. Intermediate hybrid sequences were utilized in activity screens, and they were named after the corresponding segments that differ from WT attP (e.g. for A2 attP, segment 2 in each attP half-site has been mutated to match the respective segments in Site A; Fig. S6). To predict which residues in C31-int are likely to be in close proximity to DNA, we referred to a pairwise alignment against LI integrase, for which a protein-DNA structure is available (33). Mutations made in the A2 and A4 regions most-severely inhibited wildtype C31-int (Fig. 2C-D), so these intermediates were used to screen our initial variant libraries. Regional mutagenesis was employed (codon saturation combined with insertions and deletions; also segment-specific error-prone PCR) to create amino acid diversity in stretches of the protein that were of interest. These targeted protein residues were either predicted to be close to DNA of the respective segment (34), or they were experimentally observed to be part of said interactions via pool-based alanine-knockout scans (data not shown) (35, 36).

**Figure 2:**
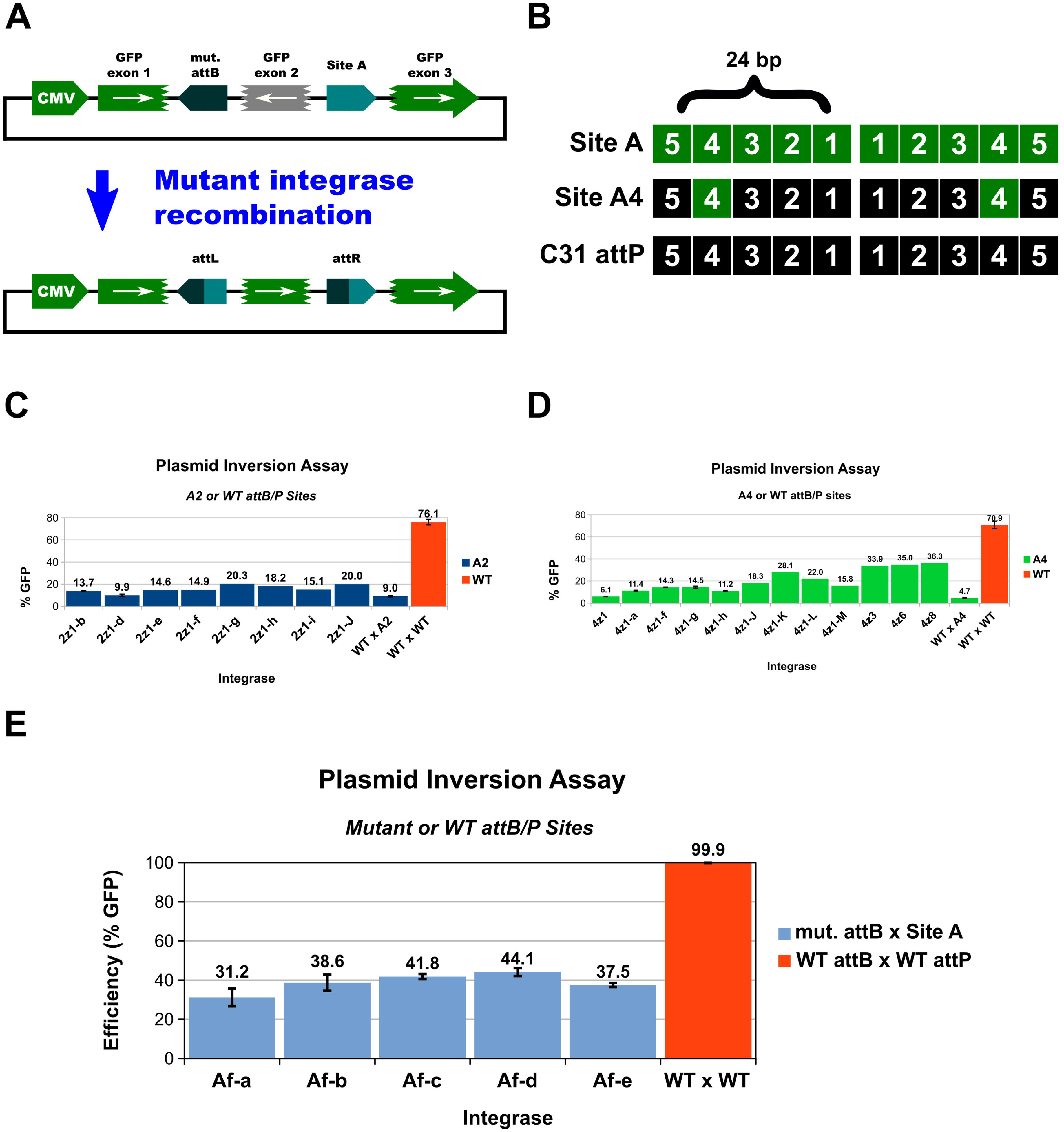
Inversion assay & screen. (**A**) To minimize false-positive signals, mutant integrases are subjected to a three-exon GFP plasmid inversion test. If no recombination occurs, the central GFP exon remains in the reverse orientation, which prevents production of green fluorescence above background. In cells with an active variant, the two attachment sites are recombined, which leads to inversion of exon 2 and the production of complete GFP. “mut. attB” refers to the attB mutant that parallels the respective attP site. E.g. when the A4 attP intermediate is used, “mut. attB” is the attB-A4 mutant. (**B**) Site A and C31 attP were divided into 5 segments, A1-A5 (1–5). Intermediate sites are named after the segment of C31 attP that has been mutated in both half sites to match the respective Site A sequence. (**C**) Variants with improved activity on A2 intermediate. WTxA2 and WTxWT show the activity of WT C31-int in the A2 and WT attP x attB inversion assays, respectively. Assay performed for 72 hours in HEK293 cells that stably express the respective variant from the H11 locus. (**D**) Variants with improved activity on A4 intermediate. WTxA4 and WTxWT show the activity of WT C31-int in the A4 and WT attP x attB inversion assays, respectively. Assay performed for 72 hours in HEK293 cells that stably express the respective variant from the H11 locus. (**E)** Five variants with the strongest ability to recombine the attachment sites of interest were tested using the plasmid inversion assay over 96 hours in HEK293 cells that stably express the respective variant from the H11 locus. For (C), (D) and (E), analysis was limited to cells that both received the inversion plasmid and that also expressed the variant integrase (single copy expressed from H11 locus). For plots with error-bars (standard error; STDEV/SQRT(3)), N=3 biological replicates.

In mammalian cells, plasmid transfection delivers hundreds-to-thousands of plasmid molecules per cell (37, 38). To maintain a 1-to-1 genotype-to-phenotype connection, we integrated a single variant per cell using an orthogonal serine integrase (Bxb1) at the H11 locus (15, 39) into a previously installed landing-pad (Fig. S4). To further support the expression of only one variant per cell, we also made use of a split-cassette system (Fig. S4), i.e. the integrated variant has a promoter upstream of its coding sequence, while non-integrated variants instead have anti-transcription elements 5’ of their ORF.

A map of the three-exon GFP recombination-reporter plasmids that we used in inversion-assay screens is shown in Fig. 2A. It incorporates an attP and attB site that flank the central inverted GFP exon – all inversion-reporter constructs had this arrangement, where each contained a specific intermediate (A1-A5) or full Site A attP-attB pair. If no recombination occurs, the central GFP exon remains in the reverse orientation, which prevents production of green fluorescence above background. In cells with an active variant, the two attachment sites are recombined, which leads to inversion of the middle exon, and thus production of complete GFP (the attL and attR sequences are spliced out with introns). FACS isolation of GFP+ cells was used to enrich active integrase variants.

All screens include an attB sequence that has been mutated in a parallel manner to the Site A attP or intermediate being tested (Fig. S6). For libraries that involve mutagenesis of the zinc-ribbon domain (ZD), which is predicted to have a major role in distinguishing attP and attB, we have mutated attB in accordance with the Rutherford et al. model, where the outermost 9 bp ZD binding segment is shifted 5 bp inward (7, 33). In libraries where the ZD has not been changed, the attB mutations simply match those made to attP (Fig. S6).

After 2-5 rounds of directed evolution – mutagenesis, activity screens, shuffling of enriched variants – we isolated a variety of A2 and A4 mutants that were able to recombine the respective intermediates at levels higher than wildtype C31-int (Fig. 2C-D). Next, we merged various A2 and A4 mutations to create “Af” (A-full) variants, and some of these were able to recombine the full Site A attP and attB sequences (top five performers shown in Fig. 2E).

### Optimization of C31-int and C31-int-dMad7 site A localization

In parallel to our directed evolution of variants, we studied optimization of wildtype (WT) C31-int localization at site A. As we are seeking to limit mutations to regions that impact DNA-interaction specificity, the ability of our variants to reach site A is unlikely to be changed from that of the wildtype enzyme. I.e., based on the LI integrase-DNA structure (33), the regions of the protein responsible for DNA interaction are likely to be separate from those that have an impact on localization. To estimate the upper limit of site A integration efficiency that we will be able to achieve, it is thus appropriate to study how well WT C31-int is able to mediate integration into an attP sequence placed at site A. In addition, any localization improvements we are able to make to WT C31-int – e.g. use of particular nuclear localization sequences (NLS), domain fusions, etc. – are likely to also improve the ability of our variants to reach site A.

To estimate WT C31-int integration efficiency at site A, we knocked-in a split-GFP recombination reporter into both loci (Fig. 3A) via Cas9-stimulated non-homologous end-joining. This system allows for measurement of integration efficiency at site A within 2-4 days, and it also supports optimization of localization efficiency via pool-based screens.

**Figure 3:**
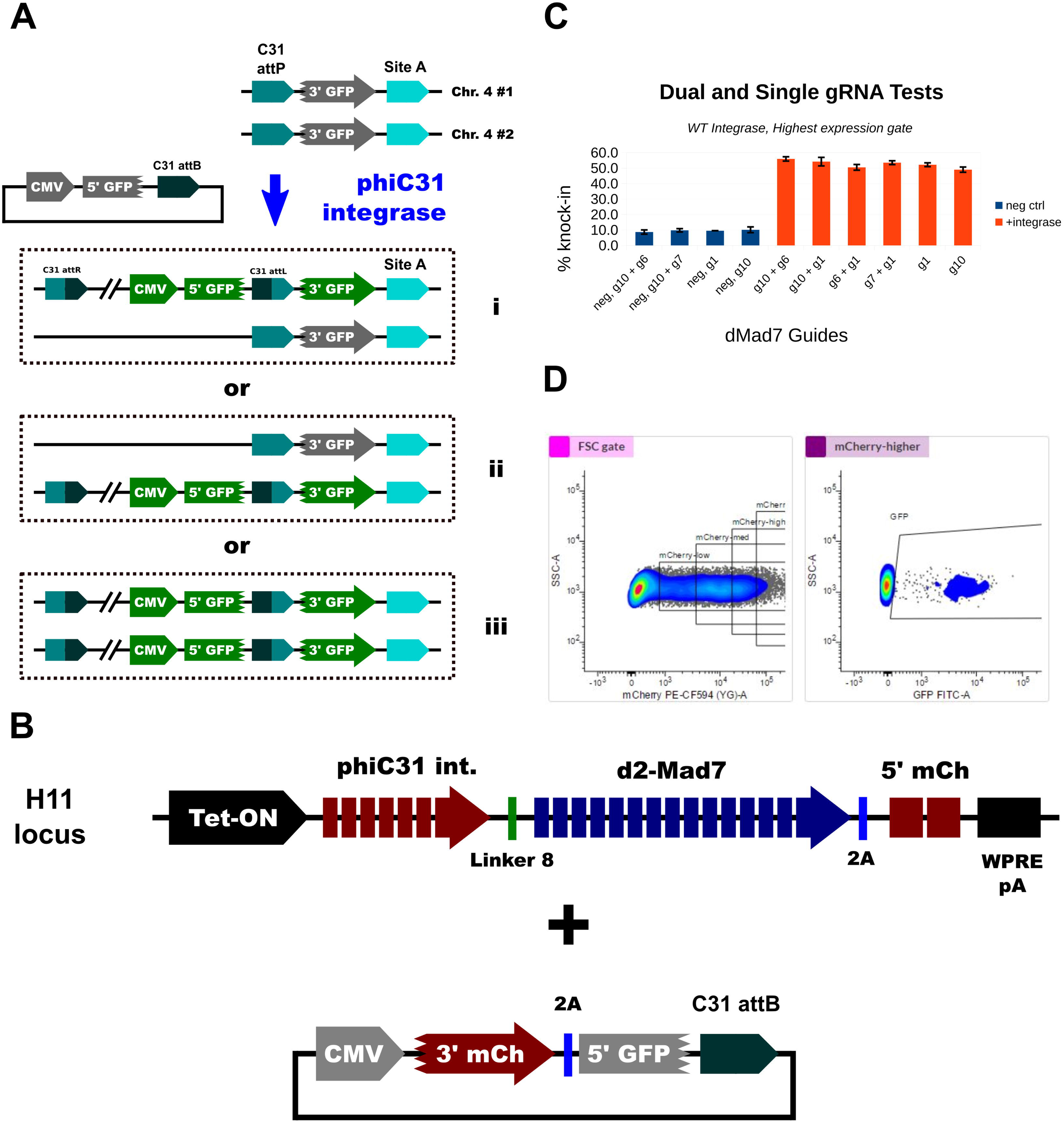
Localization optimization (**A**) Split-GFP site A integration efficiency assay. Wildtype C31 attP was placed at both site A loci in HEK293 cells. Downstream of each attP, a splice acceptor, 3’ segment of GFP, and transcription-termination sequence were also introduced. To enable detection of site-specific integration, a donor plasmid was constructed that contains the elements needed to form a complete GFP-expression cassette: CMV promoter, 5’ GFP segment, splice donor, wildtype attB site. After co-transfection of the donor and integrase-expression plasmids, cells where site-specific integration has occurred can be identified by looking for green fluorescence. Integration can happen at one (panels i and ii) or both loci (panel iii). (**B**) WT C31-int dMad7 fusion protein expression cassette used to test impact of different gRNA on site A localization efficiency. A 5’ mCherry segment fused to a trans-splicing intein domain was co-expressed and separated from int-dMad7 via a 2A-skipping peptide. In the donor plasmid, the remaining 3’ mCherry segment fused to the complementary trans-splicing intein domain was co-expressed with the 5’ GFP segment mentioned in (A), and these two proteins were separated via 2A peptide ribosome skipping. (**C**) Results of site A localization experiments for the indicated guide RNAs and combinations, in transfected cells that express the fusion protein most strongly. Assay was performed for 72 hours in HEK293 cells that stably express the fusion protein from the H11 locus. (**D**) Expression (mCherry) and GFP gating. The rightmost mCherry gate was used to analyze the cells summarized in (C).

The most impactful improvements to localization that we observed were the use of particular NLS tags, and also fusion to a deactivated Cas nuclease at the C-terminus of C31-int. A variety of groups have shown that NLS tag choice can greatly affect Cas protein performance (40–42), so to optimize this decision, we performed a screen involving 174 sequences from the N- and C-terminal regions of 87 proteins known to localize to various sub-compartments of the nucleus. The 11 most-enriched sequences were individually tested, and the best performance was observed from the C-terminal NLS of the mouse TCOF protein (UniProt O08784; data not shown).

As Cas proteins possess an active DNA-search mechanism that can penetrate chromatin, we tested the ability of C31-dCas9 and C31-dMad7 fusion proteins to improve localization efficiency. Both were able to facilitate the highest levels of integration efficiency in our split-GFP assay, however we found it necessary to heavily intronize their sequences in order to reach reasonable levels of expression (Fig. 3, Fig. S7, Fig. S3 and data not shown). Our intronized C31-dMad7 protein was expressed most strongly (Fig. S3), so we focused on this fusion approach. We tested a variety of guide RNAs that anchor the dMad7 fusion on both flanking regions of site A, and also co-expression of two guides that should anchor the protein upstream and downstream. While other choices would likely be acceptable (i.e. no statistical significance observed between guides), we focused on using the combination of guides g10 and g6 because they performed the best and have the potential to anchor integrase-dMad7 proteins on both sides of site A.

Finally, we also optimized the sequence that connects WT C31-int to dMad7. An 86-member library that mostly consisted of an NLS tag (human nucleoplasmin, mouse TCOF, or human TCOF) connected to a standard peptide linker (22 distinct sequences) was constructed and separately screened using 12 guide RNAs (Fig. S8). Two NLS-linkers dominated: NL1 and NL2 were among the top 3 most-frequently observed for 80% and 60% of the gRNA tests, respectively. We focused on these two NLS-linkers for all of our variant-dMad7 screens and tests.

### Variant-dMad7 mini-chromosome integration assay and screens

To measure variant integration at Site A in a manner similar to our localization-optimization experiments, we made several attempts to knock-in a split-GFP reporter downstream of Site A. Unfortunately, the knock-in efficiencies were too low, i.e. we could only detect the desired cells in large pools (1, 000+ cells). To create an alternative that would be compatible with pool-based screens, we implemented a SAR (Site A Reporter) mini-chromosome assay (Fig. 4A). The needed split-GFP reporter was cloned in a plasmid with 2.5 - 5 kb site A genomic sequence on each side (5-10 kb total), along with other elements needed for EBNA1-mediated extrachromosomal maintenance in human cells (EBNA1 expression, oriP, drug-resistance marker) (43). A wildtype attP version of each plasmid was also cloned for comparison to our site A split-GFP assay: we found that WT C31-int performed 5-fold worse in the SAR MAX (10 kb homology) assay, and 3-fold better in the SAR1 (5 kb homology) test (10% vs 2% and 10% vs 30%, respectively; Fig. S8A-B). We chose to focus on the SAR1 assay so that variant activity could be detected more easily.

**Figure 4:**
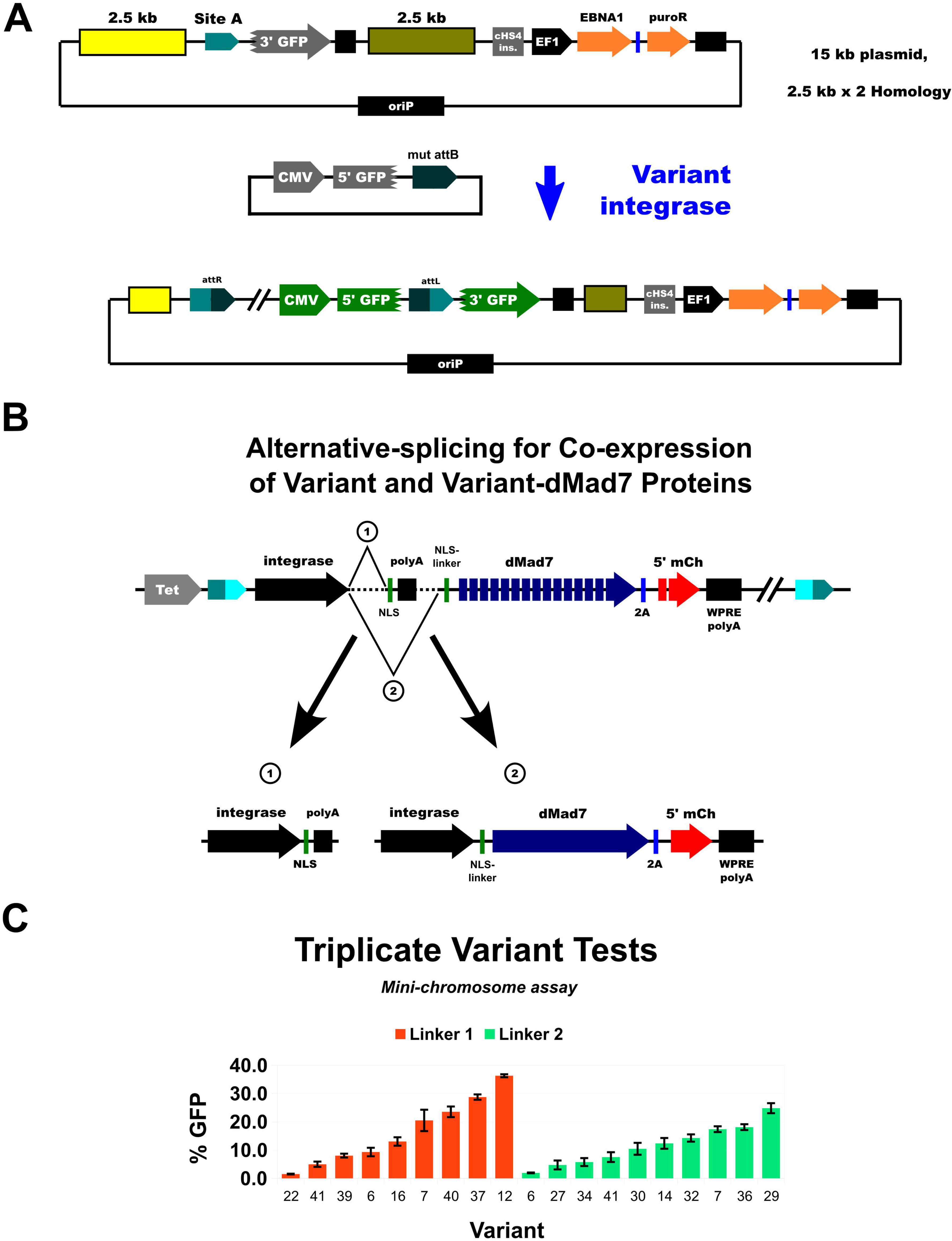
Minichromosome assay (**A**) Site A-splice acceptor-3’ GFP-polyA was cloned in a plasmid with 2.5 - 5 kb Site A genomic sequence on each side, along with other elements needed for EBNA1-mediated extrachromosomal maintenance in human cells (EBNA1 expression, oriP, puromycin-resistance marker). The 20 kb (SAR MAX, not shown) and 15 kb (SAR 1) plasmids have 5 kb and 2.5 kb genomic Site A sequence on each side (10 and 5 kb total), respectively. (**B**) When the mini-chromosome assay was used in screens, we used the depicted alternative-splicing expression construct to produce the needed solo and fused forms of each variant integrase. (**C**) Results from individual tests of the top-performing variants in the mini-chromosome assay. Assay was performed for 72 hours in HEK293 cells with the Site A SAR1 mini-chromosome that stably expressed the indicated variant using an alternative-splicing cassette from the H11 locus. N=3 biological replicates, error bars are standard error (STDEV/SQRT(3)).

When testing integrase-dMad7 fusion protein activity, we also co-express the relevant integrase protein in “solo” form without the fusion. This is done to ensure that adequate integrase dimers bind to the attB site in our donor plasmids (100s-1, 000s present per transfected cell), as the fusion proteins are difficult to express. To accomplish this in pool-based screens where a mutant attB site is used in the donor plasmid, we implemented an alternative-splicing scheme (Fig. 4B), in which each library cell produces both the solo variant protein and the variant-dMad7 fusion. To decide on the specific alternative-splicing introns to use, we tested four combinations described by Aebischer-Gumy et al. (24) in an integrase-2A-GFP/mCherry reporter (Fig. S2A and data not shown). The best performance was observed from “I4(7Y)-I4sh”, however we had to include an internal polyA sequence for optimal results (Fig. S2B). This alternative-splicing arrangement is what we implemented to produce solo variants with a C-terminal human TCOF NLS and variant-NL1/NL2-dMad7 fusions (Fig. 4B).

To identify cells that are expressing integrase-dMad7 fusions and that have also been transfected with the donor plasmid, we use a split-mCherry reporter. In the integrase-dMad7 transcript, separated by a 2A skipping peptide, we co-express a 5’ segment of mCherry fused to one half of a trans-splicing intein domain (44, 45). In our donor plasmids, the remaining 3’ segment of mCherry fused to the complementary trans-splicing intein domain is co-expressed with a 5’ segment of GFP, separated by a 2A skipping peptide. Thus, if a cell is expressing integrase-dMad7 and has also received the donor plasmid, then both halves of the mCherry protein are joined via trans-intein splicing, and red fluorescence can be produced. The 5’ and 3’ segments of GFP are seamlessly connected via RNA cis-splicing after site-specific integration.

As the mini-chromosome assay is far more challenging than plasmid inversion, we used it to screen for higher-performance variants. Two additional cycles of directed evolution were carried out using the mini-chromosome assay, and we were able to isolate a variety of mutants with improved activity from each generation (Fig. S9). We tested 9 variant-NL1-dMad7 and 10 variant-NL2-dMad7 fusions in triplicate using stable-line expression, and they had a wide range of activities (2-36%; Fig. 4C). In general, more of the top-performers came from the NL1 group (e.g. 29% for V37, and 36% for V12), however there was significant overlap (2-36% for NL1 vs 2-25% for NL2).

### Transient expression mini-chromosome assays

To move closer to real-world applications, we next tested plasmid-based transient-expression of the solo and dMad7 fusion variants. For co-expression of the two variant forms, we compared plasmids with a single alternative-splicing vector, to various ratios of vectors that separately express each protein (solo and fusion; ratios provided in supplementary materials, “Mini-chromosome establishment and assay”). For most of the variants tested, there was not a major difference between the two approaches (data not shown), so for the majority of tests we have used a single alternative-splicing plasmid to co-express solo and variant-dMad7 fusions (Fig. 5A). Overall, as expected due to the difficulty of expressing the large fusion proteins, efficiencies dropped compared to stable-line expression experiments (e.g. V12), however some variants had comparable performance (e.g. V30; Fig. 5B). Transient plasmid-based expression has the potential to temporarily express variants at higher levels than is possible from single-copy stable lines, so we also tested weaker expression of certain variants (i.e. mPGK promoter instead of CAG; Fig. 5C). For certain variants, this led to efficiencies that were competitive with our top strong-promoter reference (CAG-V12, ratio E), so clearly there is no single optimal expression level for our variants.

**Figure 5:**
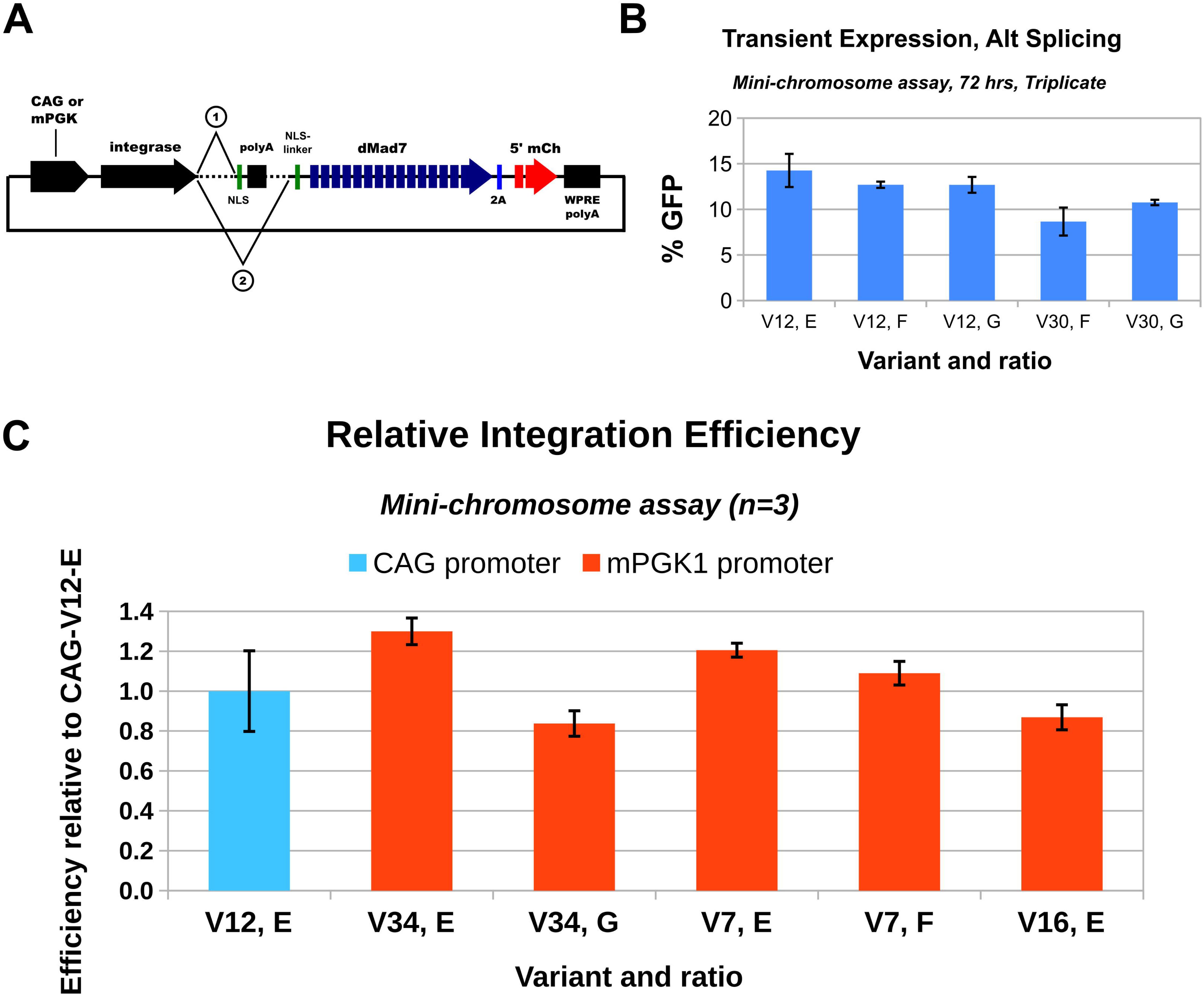
Transient variant expression, mini-chromosome assay (**A**) Drawing of the plasmid used to express solo and fused forms of variants via alternative-splicing in transient-delivery assays. hU6-driven expression cassettes for guide RNAs g10 and g6 were also present in these vectors (not shown). (**B**) Results from V12 (ratios E, F, G) and V30 (ratios F, G) expressed via the CAG-promoter in the transient expression mini-chromosome assay. (**C**) Results for mPGK-promoter driven expression of V34 (ratios E, G), V7 (ratios E, F) and V16 (ratio E), normalized to CAG-driven V12 ratio E. All experiments were performed over 72 hours in HEK293 cells that contained the mini-chromosome (Site A SAR1). Ratios have been provided in the “Mini-chromosome establishment and assay” section of the supplementary materials. N=3 biological replicates and error-bars are standard error.

### Stable line and transient expression native site A integration assays

Our final tests estimate the integration efficiency of variants at unmodified (native) site A. The 10.2 kb donor plasmid co-expresses full GFP, 3’ mCherry fused to a trans-splicing intein domain, and a puromycin-resistance marker, all separated by 2A skipping peptides (Fig. 6A). For integration-efficiency measurement, a barcode pool (18 nucleotide G/T; K18) was cloned near the mutant attB site. Specific amounts of transfected cells were isolated via FACS three days after plasmid delivery, and they were then outgrown in medium supplemented with puromycin (to remove cells that haven’t stably integrated the plasmid). To estimate targeted integration efficiency, which is the number of observed unique barcodes divided by the sorted cell count, a region encompassing the barcode segment and integration junction was amplified via nested genomic PCR and then sequenced (Fig. 6B). We use the term “efficiency” because a known amount of cells were sorted before drug-selection was started, i.e. this is not a measurement of specificity, and no off-target analysis has been performed. When this assay was performed with HEK293 cells stably-expressing V12-NL1a4-dMad7, V37-NL1a4-dMad7 or V30-NL2-dMad7, we observed 32.4%, 6.94% and 15.5% percent integration efficiencies, respectively (Fig. 6C). For transient plasmid-based expression, only the mPGK-V7-NL2a4-dMad7 vector yielded detectable integration efficiencies, and the values ranged from 2.1% - 13.3% (Fig. 6C). Ratios E and F yielded integration efficiencies close to what was observed for V7 in the mini-chromosome assay, however the conditions were quite unforgiving, i.e. small changes in confluency at the time of transfection majorly impacted the outcomes (e.g. for ratio E, 2.8% vs 6.2-13.3% for a Higher Cell Density (HCD); visa-versa for ratio F).

**Figure 6:**
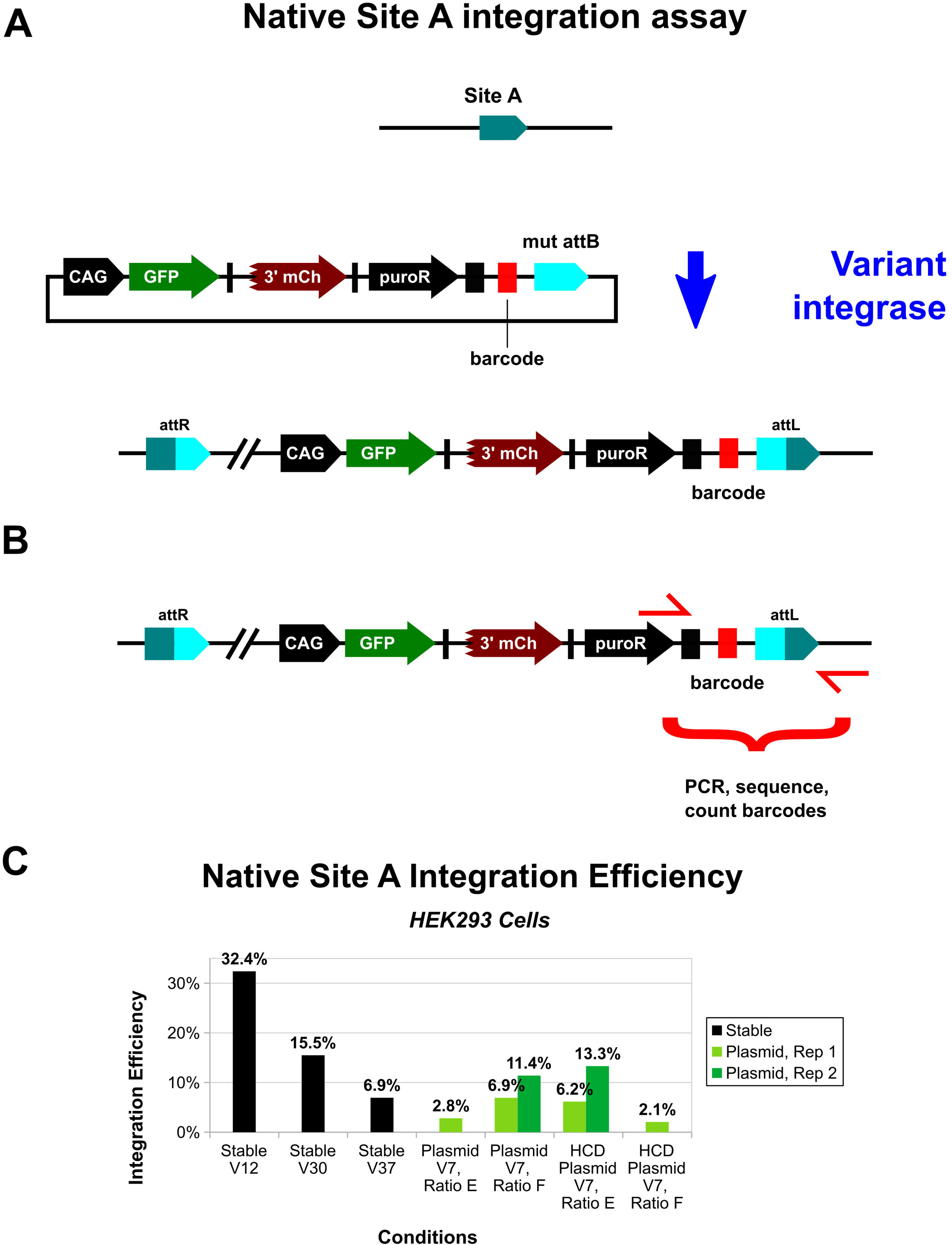
Native site A assay (**A**) The 10.2 kb donor plasmid co-expresses full GFP, 3’ mCherry fused to a trans-splicing intein domain, and a puromycin-resistance marker, all separated by 2A skipping peptides. A barcode pool (18 nt G/T; K18) was cloned near the mutant attB site. (**B**) Specific amounts of transfected cells were isolated via FACS three days after plasmid delivery, and they were then outgrown in medium supplemented with puromycin. To estimate integration efficiencies, the depicted region encompassing the barcode segment and integration junction was amplified via nested genomic PCR and sequenced, then barcodes were counted. (**C**) Barcode-counting results for stable and transient-expression Native Site A assay. For Stable V12, V30 and V37, the expression constructs were V12-NL1a4-dMad7, V30-NL2-dMad7 and V37-NL1a4-dMad7, respectively. NL1a4 refers to alternative splicing with NLS-linker-1. V30-NL2-dMad7 was not expressed using alternative splicing, so a separate solo V30 expression plasmid was co-transfected with the donor. For transient plasmid-based expression, only the mPGK-V7-NL2a4-dMad7 vector (ratios E and F) yielded detectable integration efficiencies. HCD: Higher Cell Density.

## DISCUSSION

In this study we have described the development and initial characterization of S-SELeCT integrases that have been evolved to target a novel safe-harbor site in human locus 4p14 (Site A). To our knowledge, this is the first time that a serine integrase has been modified to act on an endogenous symmetrical attachment site (att-site) in the “landing-pad” mode, i.e. where integrase homodimers carry-out the recombination reaction to completion. This is in contrast to pseudosite integration (using homodimers), where assistance from host DNA-repair machinery is often needed, and can result in small indels, large deletions and inversions.

It is also our understanding that we are the first to conduct pool-based activity screens in mammalian cells as part of the directed evolution process for altering serine integrase specificity (Fig. 7). This is slower than screening in prokaryotes, however it was necessary in our case because C31-int-family variant activity observed in bacteria has often not translated into mammalian cells (17, 18).

**Figure 7:**
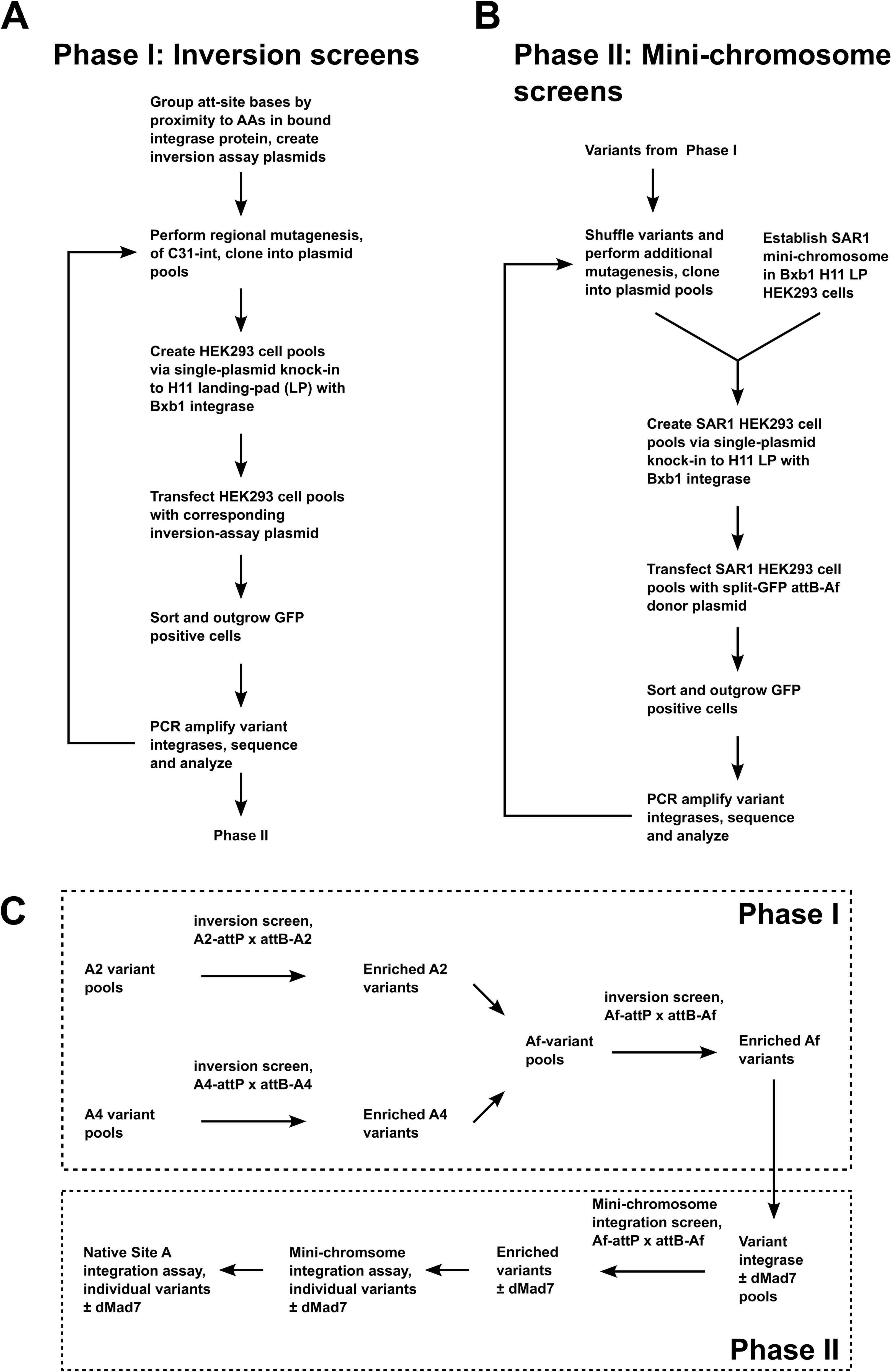
Workflow for inversion-reporter and mini-chromosome-assay screens (**A**) Overview of phase I cycles. In phase I, we screened variants using inversion assay screens. (**B**) Overview of phase II cycles. In phase II, we took variant sequences obtained from phase I, performed additional mutagenesis, then screened using the mini-chromosome assay. (**C**) Complete overview. Inversion screens started in parallel using A2-attP x attB-A2 and A4-attP x attB-A4 recombination challenges. Active variants enriched from multiple rounds of these screens were then combined to create Af-variant pools, which were screened using Af-attP (50 bp Site A full) x attB-Af (40 bp attB-A full) inversion assays. In phase II, enriched variants from these pools were screened using the mini-chromosome assay. Alternative splicing was used in these phase II screens to co-produce solo and dMad7-fusion forms. Individual variants obtained from enriched pools were then subjected to testing in the mini-chromosome and native Site A assays.

While pool-based mammalian screens uniquely enable high-throughput optimization of expression, nuclear localization, and fusion-protein linkers, they also critically minimize prokaryotic dead-end-variants. I.e. the poor performance of previous variants cannot be blamed on expression or localization problems because they shared these features with their wildtype counterparts (e.g. C31-TG1 hybrids not active in mammalian cells, but both parental integrases were) (18). In addition, mammalian screens implicitly select against sequences that receive inhibitory post-translational modifications and that have problematic interactions with host cell proteins. Mammalian libraries have been more difficult and time consuming to establish because unlike bacteria, plasmid delivery (transfection) is not clonal (instead of single plasmid delivery, 100s-1000s are introduced per cell). Serine integrases and pool-based screening methods like those we have described here offer the most efficient solution to these problems, i.e. they are the best at performing large-insert site-specific single-copy integration (46), and this enables the reasonable establishment of libraries with diversities in the 1e6-1e7 range and higher (e.g. 1e8-1e9 with large-volume or flow electroporators).

To optimize the ability of variants to reach Site A, we created a reporter line where an attP-3’GFP-polyA segment was integrated into both copies of the locus (Fig. 3A). This line was straightforward to create and proved to be quite useful, so we made several attempts to create a version that could be used to detect site A integration. Unfortunately, SpCas9 and Mad7 targets in desirable regions outside of site A were far less efficient at stimulating knock-in of our 3’GFP-pA insert, so we opted to instead use an EBNA1-oriP extra-chromosomal plasmid (mini-chromosome) where the reporter segment was flanked by site A genomic sequence (5.0 kb total, 2.5 kb on each side; Fig. 4A). This system was used successfully to (i) optimize NLS-tag-linker sequences, (ii) select helpful dMad7 guide RNAs, and (iii) isolate variants that can mediate integration into Site A. The mini-chromosomal system was not always predictive of native Site A integration efficiency, however they are conducted on different time scales (3 days vs 1-2 weeks), so phenomena that take longer to manifest could be at least partially to blame. E.g. efficiencies might drop for variants that have toxic side-effects, and conversely efficiencies could increase for variants that act more slowly (we have observed the latter for certain variants, e.g. Fig. S10). Disruption of C31-int unidirectionality, i.e. creation of mutants that perform attL x attR recombination without the RDF, could also lead to a disconnect between the two assays, however we avoided mutagenesis of the coiled-coil (CC) region that controls this (as shown in tables S1-S8, our mutation positions range 114-391; CC spans 452-518) (47).

As we anticipated, transient expression via plasmid delivery majorly reduced integration efficiency (Fig. 4C vs Fig. 5B-C). We were still able to measure integration efficiency for V7, however only when we used a weaker promoter (mPGK) than in the initial tests (CAG), and there was major sensitivity with regards to the confluency of the cells on the day of transfection (which is difficult to control and measure; Fig. 6C). Screens were performed using single-copy expression, so for more optimal results, it might be necessary to implement more sophisticated expression control (to ensure an even and low-level in more cells). It is likely that RNP and/or mRNA would improve results and consistency even further, as they have for a variety of Cas-based gene-editing approaches. However, these are not as trivial to produce given the prokaryotic disconnect observed for C31-int-family proteins (e.g. protein might not fold the same) and the large sequences involved (e.g. truncated mRNA could poison integrase tetramers). Smaller fusion partners would make plasmid, RNP and mRNA delivery more tractable, so they are a worthy future direction to consider.

For native Site A efficiency measurement, we focused on barcode counting to avoid potential pitfalls that can be encountered with ddPCR and other quantitative PCR methods. For example, Pandey et al. (48) recently pointed out a major flaw in the primers used to measure PASTE efficiencies, which may have at least partially contributed to their over-estimations. ddPCR can be used to accurately estimate efficiencies, but calibration with appropriate controls is critical for accuracy (and barcoding can help with that; (48)).

Barcode counting is not free of problems, and we encountered several, including major PCR skew in barcode counts (Fig. S11), PCR noise, and sequencing miscalls. In the future, to reduce skew, different donor backbones and primers should be tested to improve amplification efficiency. If this is successful (e.g. nested PCR no longer needed), then emulsion or droplet PCR could be considered for further reduction of over-representation (i.e. by barcodes that get amplified early in the reaction) (49).

With regards to alternative gene-editing approaches, we consider ee/evoPASSIGE and MINT to be the most similar to ours (12, 48). While the PASSIGE methods allow for rapid development of targeting systems, they are all based on Prime editing, which involves a series of enzymatic steps (nicking of multiple DNA strands, reverse transcription, flap resolution) that can generate undesired outcomes like indels at off-target sites (48, 50, 51). For example, in a recent analysis, Witte et al. point out that eePASSIGE targeted at AAVS1 was reported to have 1.6X more off-targets than on-targets (see Table S6 of Science publication, not PMC version), which was similar to their modified transposon system (48, 52).

Sangamo’s MINT system utilizes serine integrases with modified specificity like we do, however they are focused on integrating into endogenous asymmetric att-sites (12). While carrying out directed evolution screens, their process is conducted in bacteria: full and intermediate symmetric att-sites based on each half of the asymmetric endogenous site are used. Then, to test integration in human cells, they combine a top-performer from each half-site screen with the intention of forming heterodimers that will mediate integration into the endogenous asymmetric site.

Integrating into asymmetric sites that are not necessarily pseudosites vastly increases the number of potential genomic locations that can be targeted (i.e. the only sequence-based requirements would be uniqueness and a non-palindromic di-nucleotide core). However, forcing heterodimerization of serine recombinase catalytic domains would not be trivial and has the potential to create undesirable DNA damage, so this approach is likely to always have a more limited integration-efficiency ceiling compared to symmetric-att-site integration. This is because each integrase variant can form homodimers, which limits the amount of desired tetramers in cells. MINT also uses the WT integrase, which further limits integration efficiencies. This is because WT Bxb1 int can heterodimerize with each variant, and none of those are productive for the desired reaction. In the longer-term, tyrosine integrases would be a better choice for asymmetric integration because the reaction mechanism would not be in conflict with forced heterodimerization (and there is precedence for this with Cre (53)), however the problem currently is that they are not nearly as efficient as high-performance serine integrases (e.g. Voziyanova et al.) (54).

For S-SELeCT, additional important future directions include (i) more rounds of directed evolution, (ii) development of variants that target different safe-harbor sites, (iii) integration efficiency measurements in other cell types such as primary cells and iPSC, (iv) off-target analysis.

Additional rounds of directed evolution are warranted because we observed improvements in efficiency for previous rounds (Fig. S9), and we have not yet reached the maximum that was observed in our WT C31-int experiments (∼50%; Fig. 3C) -- i.e. there is clearly room to grow.

Variants that target different locations should be developed because site A is a novel safe-harbor locus that might not prove ideal for certain therapeutic applications. E.g. integrations into this locus could trigger unwanted impacts on the expression of various proteins/ncRNA, or expression might be silenced after differentiation into particular lineages, or the locus could be unavailable for integration in certain cell types (either directly because of heterochromatin or indirectly because of inhibitory proteins).

Integration efficiency measurements and off-target analysis should be performed in potential primary cells of interest because these outcomes cannot be predicted, and immortal polyploid lines like HEK293 are not representative. Still, we did not make use of HEK293 cells that express the SV40 T antigens, so in combination with RNP-delivery, 10-30% integration efficiencies in permissive primary cells (e.g. iPSCs) of a 10 kb plasmid would not be unreasonable. We expect that even larger cargo targeting will be possible, however at lower efficiencies, because this is a well-established observation for robust serine integrases (including phiC31), and we have not modified regions of the protein that are involved in the reaction fundamentals (monomer dimerization, dimer synapsis, strand-exchange, etc.) (55–59). We’re also doubtful that off-target integrations are a major occurrence for the current generation of S-SELeCT integrases fused to dMad7, however they very likely will be in future derivative variants that have higher efficiencies, as is often the case for a variety of serine integrases and other gene-editing systems. We have this opinion for four major reasons: (i) parts of the integrase that are likely to create hyperactivity were not mutated (e.g. as was done for ee/evoPASSIGE (48)), (ii) our assays only yield signal when sequence-specific recombination has occurred, (iii) the nature of the mutations we’ve observed to date do not point to specificity-reduction (e.g. very few conversions to alanine, glycine, etc.; Tables S1-S8), and (iv) no results from inversion-assay experiments have supported specificity loss. If we were weakening specificity, then variants would have predominantly interacted with DNA elsewhere in the cell instead of our plasmids, thus decreasing GFP signal in the assays. This is a phenomenon that was observed when studying the phiC31 RDF, i.e. the pseudosite-prone pCSI integrase expression vector mediated less attP x attB recombination than the pCS-kRI plasmid (which co-expresses the inhibitory RDF), so pCS-kI was created (pCS-kRI without RDF) and it was observed to perform far better at WT recombination (15, 60) (and data not shown). In addition, reduced specificity would not have produced outcomes like we observed for variants 4z-1 and 4z-2, which have majorly lost the ability to recombine WT att-sites, but handle the A4 intermediates better (4z-1) or at a similar low rate (4z-2) (Fig. S12).

## AUTHOR CONTRIBUTIONS

Alfonso Farruggio: Conceptualization, Data curation, Formal analysis, Funding acquisition, Investigation, Methodology, Project administration, Software, Supervision, Validation, Visualization, Writing – original draft, Writing – review & editing. Lin Jiang: Data curation, Investigation, Supervision, Validation, Writing – review & editing. Karen Duong: Data curation, Investigation, Supervision, Writing – review & editing. Cynthia Nguyen: Investigation. Razan Kaddoura: Data curation, Investigation, Supervision. Ruby Tsai: Conceptualization, Funding acquisition, Project administration, Supervision, Writing – review & editing.

## SUPPLEMENTARY DATA

Supplementary Data are available at NAR online.

## DATA AVAILABILITY

The data underlying this article are available in the article, in its online supplementary material, and in Zenodo at https://doi.org/10.5281/zenodo.18356960.

