## Supplementary Figures for "S-SELeCT: A Human-Evolved Serine Integrase System for Efficient Large-Cargo Genome Integration"

**Figure S1: Comparison of wildtype Mad7 and two candidates for deactivated Mad7 in the EGxxGFP activation assay**

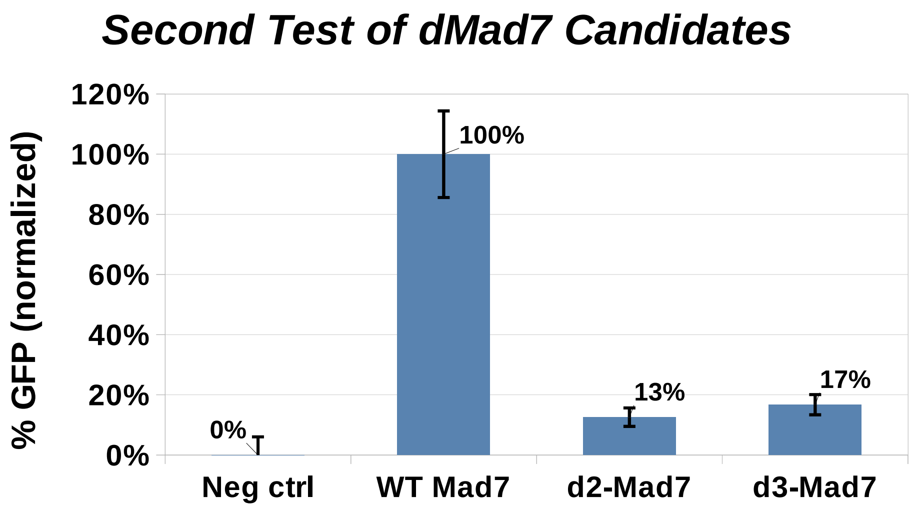


In the EGxxGFP assay, EGFP signal is produced if homologous recombination is used by the host cell to repair the transfected reporter plasmid. A target sequence for Mad7 nuclease is flanked by homologous EGFP sequence, so if cleavage of the target stimulates homologous repair, then the coding sequence of EGFP is produced. Otherwise, the target and flanking homology prevent the formation of EGFP, i.e. the coding sequence is interrupted. d2-Mad7 (called “dMad7” in this manuscript) is D877A, E962A. d3-Mad7 is D877A, E962A, D1213A.

**Figure S2:** Assay used to compare alternative-splicing introns

A

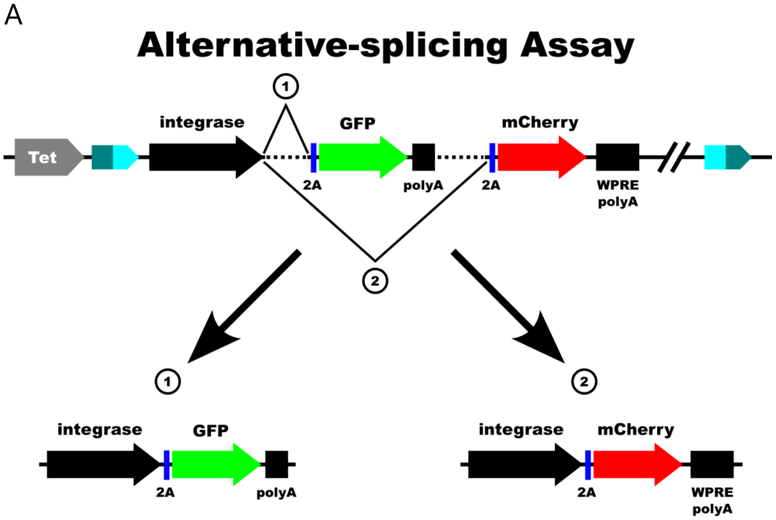


B

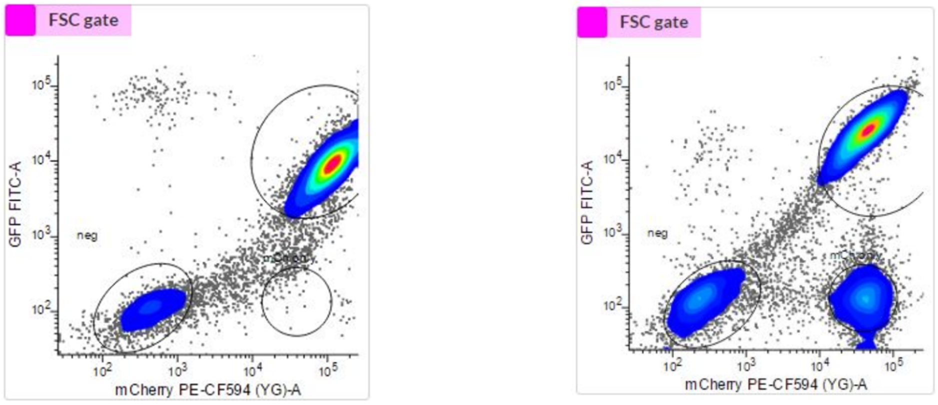


**With** internal polyA Without internal polyA

(A) The depicted expression cassette, and also a no-internal-polyA version with no polyA 3′ of GFP (not shown), were used to test four combinations of alternative-splicing introns from Aebischer-Gumy et al. 2023: I4(7Y)-I4sh, I4(3Y)I4sh, I4(5Y-5)I4, I4(0Y)I4. If the first splice occurs (1), then WT C31-int is fused to 2A-GFP-polyA, or to 2A-GFP with 2A-mCherry as part of the 3′UTR in the no-internal-polyA version. If instead the second splice happens (2), then then WT C31-int is fused to 2A-mCherry-WPRE-polyA. (B) Comparison of the I4(7Y)-I4sh combination of introns with and without the internal polyA. Vertical axis: GFP; horizontal axis: mCherry.

**Figure S3:** Comparison of transient expression strength of integrase-dCas cassettes

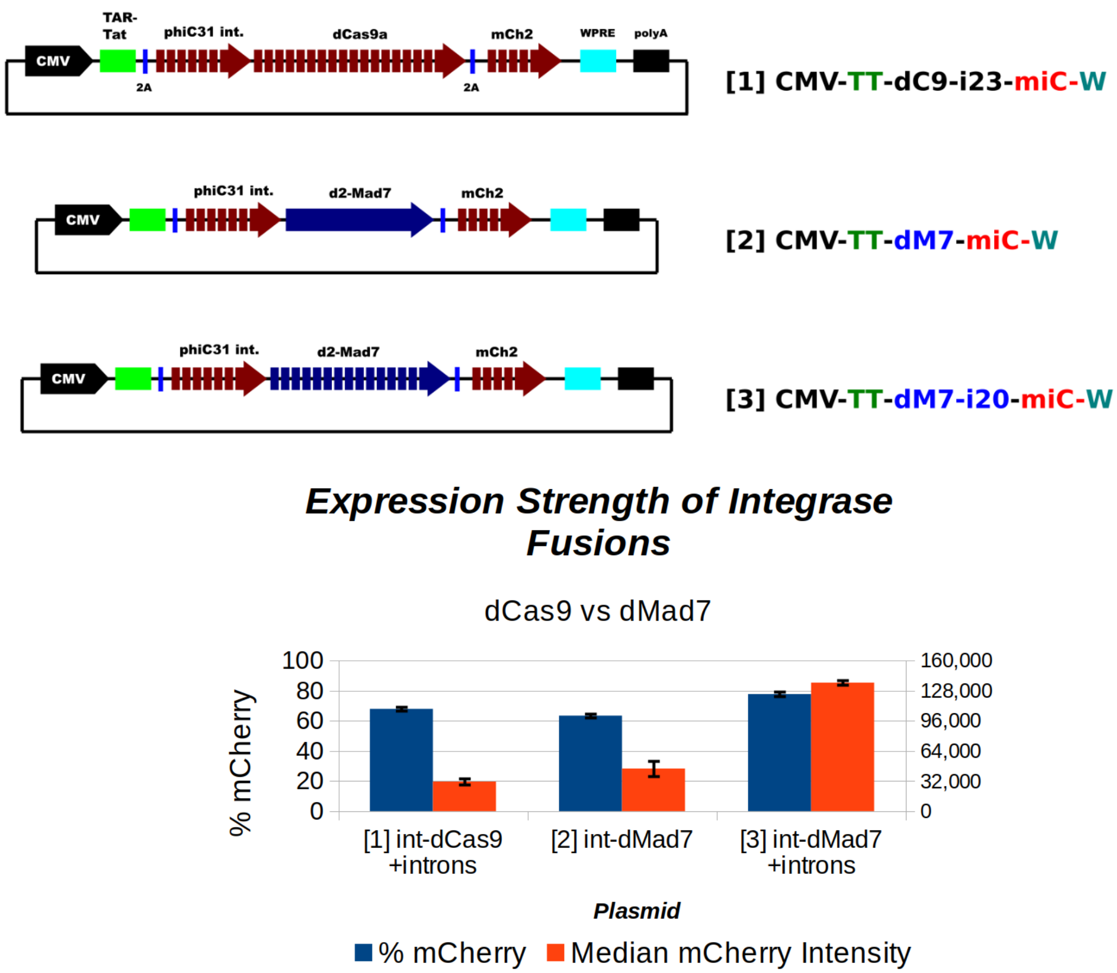


Top: maps of the expression plasmids tested. All expression cassettes were driven by the CMV promoter, have a 5′ HIV TAR-Tat sequence (TAR sequence-kozak-Tat ORF-2A), 3′ 2A-intronized mCherry, 3′ WPRE and a 3′ BGH UTR/polyA. [1] intronized integrase-dCas9a. [2] intronized integrase fused to dMad7 with no introns. [3] intronized integrase-dMad7. Bottom: HEK293 cells were transfected with each plasmid, and then analyzed for mCherry expression after 72 hours. The blue bars show the average percent of cells that were positive for mCherry (left axis), while the red bars show the median mCherry intensity (right axis). N=3 biological replicates, error-bars are standard error.

**Figure S4:** Mammalian stable-line expression

**A**

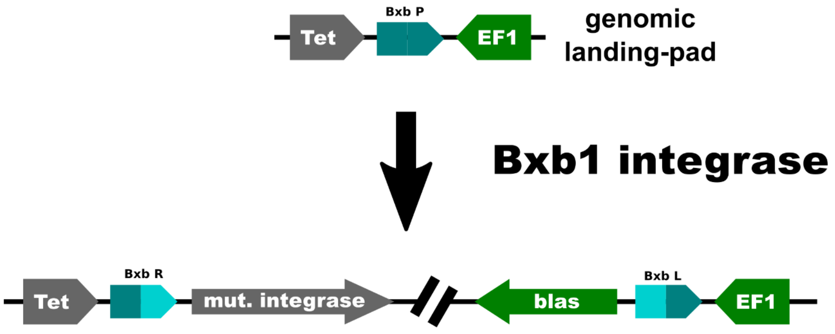


**B**

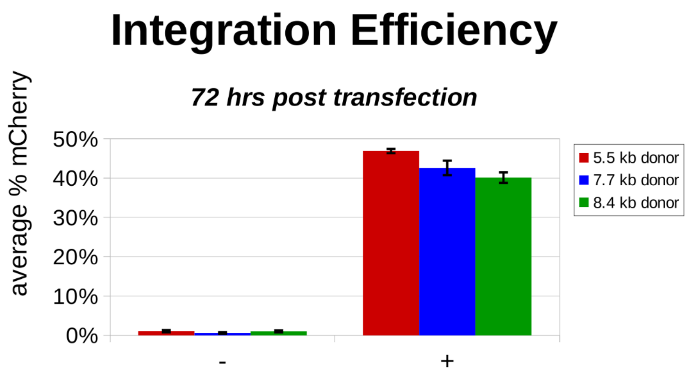


(A) A Bxb1 attP (Bxb1 P) landing-pad was placed at the H11 locus in HEK293 cells that had been previously engineered to express the rtTA-V10 transcription factor [Zhou 2006]. As pictured, the Tet [Loew 2010] and EF1alpha (EF1) promoters are oriented in a convergent manner towards the attP site. Site-specific integration is mediated by co-transfection of a Bxb1 integrase expression vector with the donor plasmid (not shown). After site-specific integration of the donor, the EF1 promoter in the landing-pad drives marker expression (e.g. blasticidin or puromycin-resistance, mCherry, etc.). Tet is activated via doxycycline supplementation when expression of integrase proteins is desired. (B) To measure integration efficiency, various donors that only express mCherry after site-specific integration were co-transfected with a Bxb1 integrase expression plasmid, and after 72 hours red fluorescence levels were measured. For negative controls, each donor was instead co-transfected with a filler plasmid (same backbone as Bxb1 integrase expression vector). N=3 biological replicates, error-bars are standard error.

**Figure S5:** Genomic context of Sites A-i and Xq22

| site code name | Locus | 5’ gene | 5’ gene distance [kb] | 3’ gene | 3’ gene distance [kb] |
|---|---|---|---|---|---|
| A | 4p14 | LINC01258 | 65 | KLF3-AS1 | **20** |
| B | 6p21.2 | MOCS1 | 385 | LINC00951 | **20** |
| C | 9q34.3 | FCN1 | 120 | **OLFM1** | **35** |
| D | 11p12b | C11orf74 | 1085 | LOC105376633 | 190 |
| E | 11q22.3 | C11orf87 | 180 | ZC3H12C | 480 |
| F | 14q21.2 | MIS18BP1 | **14** | LINC00871 | 795 |
| G | 1q41 | LOC102723886 | **6.5** | SLC30A10 | **350** |
| H | 11p12a | C11orf74 | **1045** | LOC105376633 | **230** |
| i | 12q24.32 | LOC101927592 | **115** | LOC101927616 | 145 |
| Xq22 | Xq22.1 | CENPI | 30 | DRP2 | 20 |

Identified attP-like sites, including A – i, and a previously reported pseudo site Xq22.1. For each site, the closest annotated 5′ and 3′ genes are listed, along with their distance.

**Figure S6:** Sequences of intermediate and full att-sites

| | |
|---|---|
| A full | 5'-**GAGCATCCCCAACGAAGAGGACTT TC AGGTCTCCTTATTTGGGAAAACCC**-3' |
| A1 | 5'-gtagtgccccaactggggtaac**TT** TT **AG**gttctctcagttgggggcgtag-3' |
| A2 | 5'-gtagtgccccaactggg**AGG**acct TT gagt**CTC**ctcagttgggggcgtag-3' |
| A3 | 5'-gtagtgccccaa**CGAA**ggtaacct TT gagttctc**TTAT**ttgggggcgtag-3' |
| A4 | 5'-gta**CATC**cccaactggggtaacct TT gagttctctcagttggg**AAAA**tag-3' |
| A5 | 5'-**GAGC**tgccccaactggggtaacct TT gagttctctcagttgggggc**ACCC**-3' |
| C31 attP | 5'-gtagtgccccaactggggtaacct TT gagttctctcagttgggggcgtag-3' |

| | |
|---|---|
| attB-A full | 5’-**GAGCATCCCAAGAGGACTT TC AGGTCTCCTTAGAAAACCC**-3’ |
| attB-A1 | 5’-cgggtgccagggcgtgc**TT** TT **AG**gctccccagggcgcgta-3’ |
| attB-A2 | 5’-cgggtgccaggg**AGG**gccc TT gggc**CTC**ccagggcgcgta-3’ |
| attB-A3 | 5’-cgggtgc**CCAA**gcgtgccc TT gggctccc**TTAG**gcgcgta-3’ |
| attB-A4 | 5’-cgg**CATC**cagggcgtgccc TT gggctccccagg**AAAA**gta-3’ |
| attB-A5 | 5’-**GAGC**tgccagggcgtgccc TT gggctccccagggcg**ACCC**-3’ |
| C31 attB | 5'-cgggtgccagggcgtgccc TT gggctccccagggcgcgta-3' |

Segment A1-A5 intermediate attP and attB sequences. For clarity, the complete sub-sequences from Site A are bolded, capitalized and colored, even if particular bases match in the WT C31 position. The Site A sub-sequences for A1, A2, A4 and A5 are identical in attP and attB. For attB-A3, the optimal sub-sequences were determined via a pool-based screen.

**Figure S7:** Comparison of stable line expression of C31-int-dCas9a fusions without and with introns

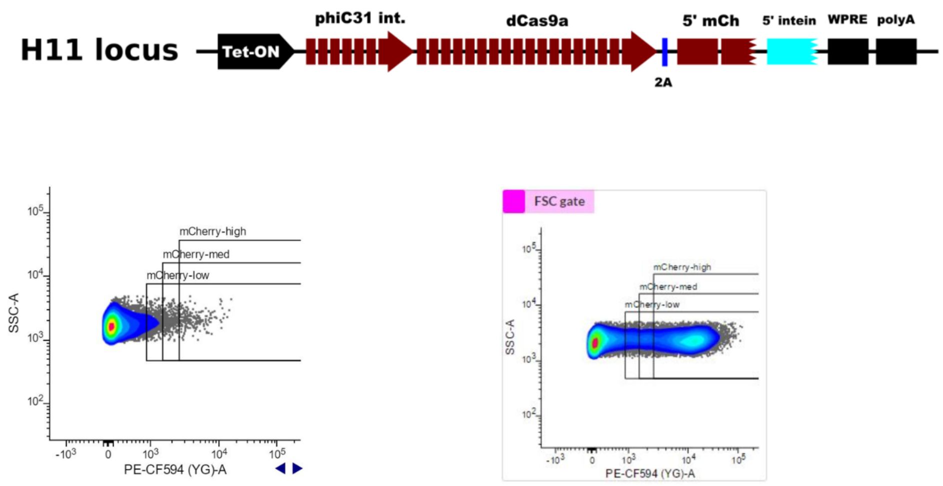


**Stable C31-dCas9-mCherry,**

**No introns**

**Stable C31-dCas9-5' mCherry-5' intein,**

**Heavily intronized**

Top: Expression cassette used to express heavily-intronized integrase-dCas9a-2A-5'mCherry-5'intein transcript. Bottom left: flow-cytometry plot of mCherry fluorescence observed when expressing a no-intron integrase-dCas9a-2A-mCherry transcript. Bottom right: mCherry fluorescence when the depicted cassette was expressed in cells that were also transfected with a plasmid that expresses 3'intein-3'mCherry. In both cases, expression occurred from the H11 locus in HEK293 cells, and the cells were incubated in doxycycline for 96 hours. The 3'intein-3'mCherry plasmid was transfected 24 hours after initiation of Tet expression (i.e. 72 hours before the data was collected).

**Figure S8:** NLS-Linker screen.

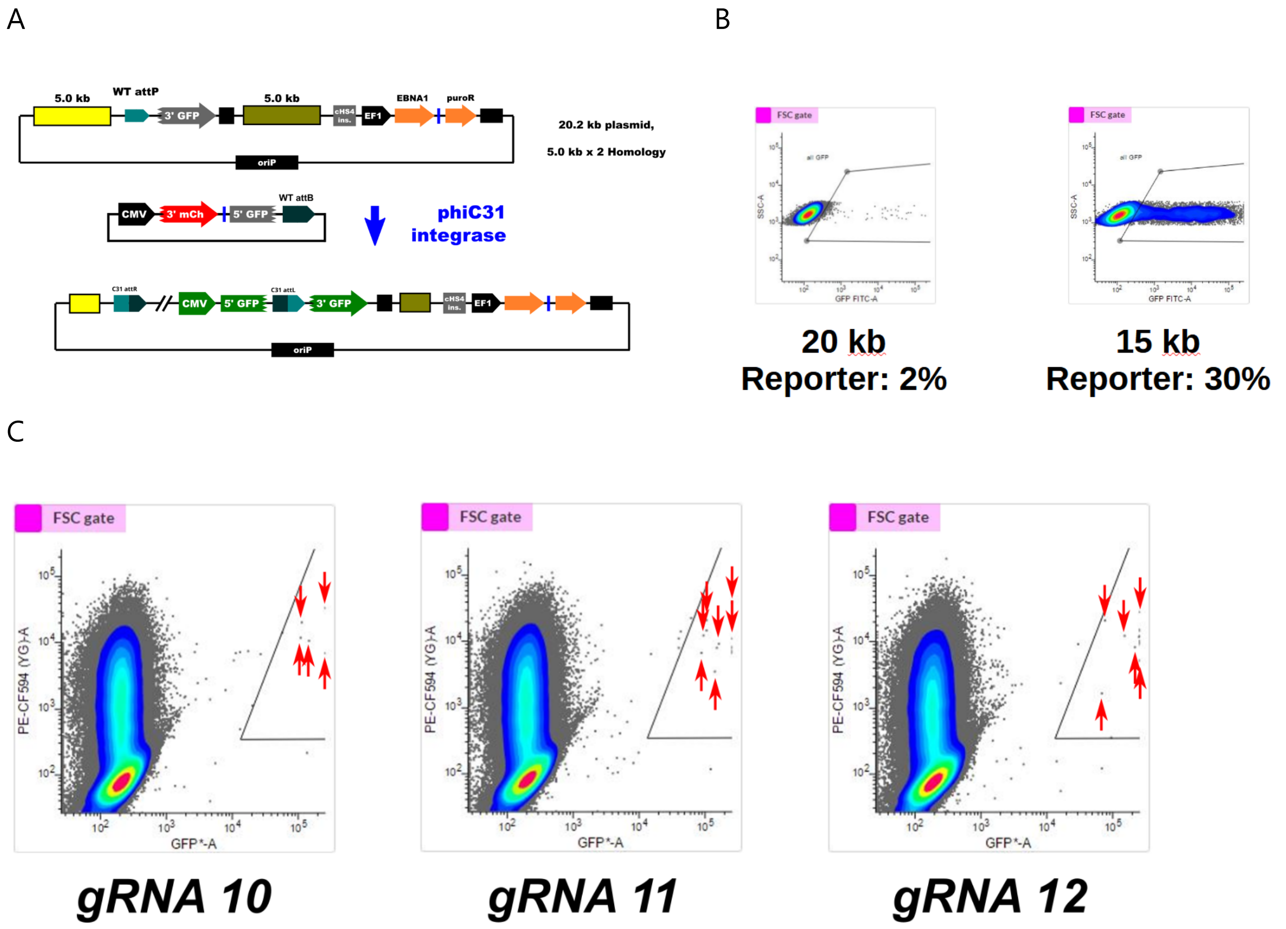


(**A**) Wildtype att-site SAR MAX Mini-chromosome assay used to screen NLS-linker libraries. mCherry signal is produced in transfected cells that also express the WT-C31-dMad7 fusion via trans-intein splicing. 5′ mCherry-5′intein is co-expressed in the WT-C31-dMad7 transcript and separated via 2A skipping peptide (not shown). 3′intein-3′mCherry-2A-5′GFP is expressed via the donor plasmid as shown. (**B**) Both wildtype att-site SAR plasmids were tested to compare C31-int performance to what we observed in our Site A split-GFP assay (Fig. 3a). We found that WT C31-int performed 5-fold worse in the "SAR MAX" assay, and 3-fold better in the "SAR1" test (10% vs 2% and 10% vs 30%, respectively). For higher-stringency, the MAX WT att-site version (20 kb plasmid) was used in NLS-linker screens. (**C**) WT C31-int fusions to dMad7 were screened in pools, and those that produced sufficient GFP signal (horizontal axis) relative to the level of expression (mCherry; vertical axis) were sorted. The gates used for this and three examples collected during sorts are shown. Red arrows point to various positive events that were sorted.

**Figure S9:** Generational activity improvement

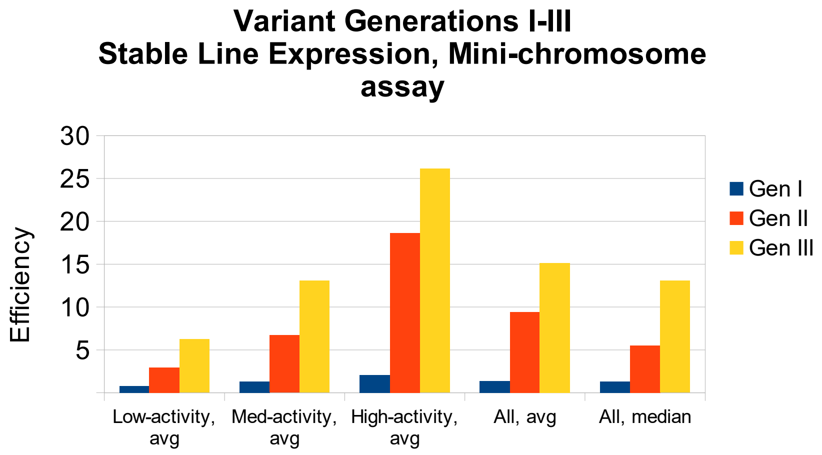


Variants from three successive generations (I, II, III) of directed evolution were separated into low, medium and high-activity groups. Within each generation, the three groups have equal sizes (e.g. Gen II low, med and high all have 8 members). The average activity level (integration efficiency in mini-chromosome assay) of each grouping is shown. Also shown are the average and median activity levels for the three generations with no sub-groups ("All, avg" and "All, median", respectively).

**Figure S10:** Longer-term tests of variant activity.

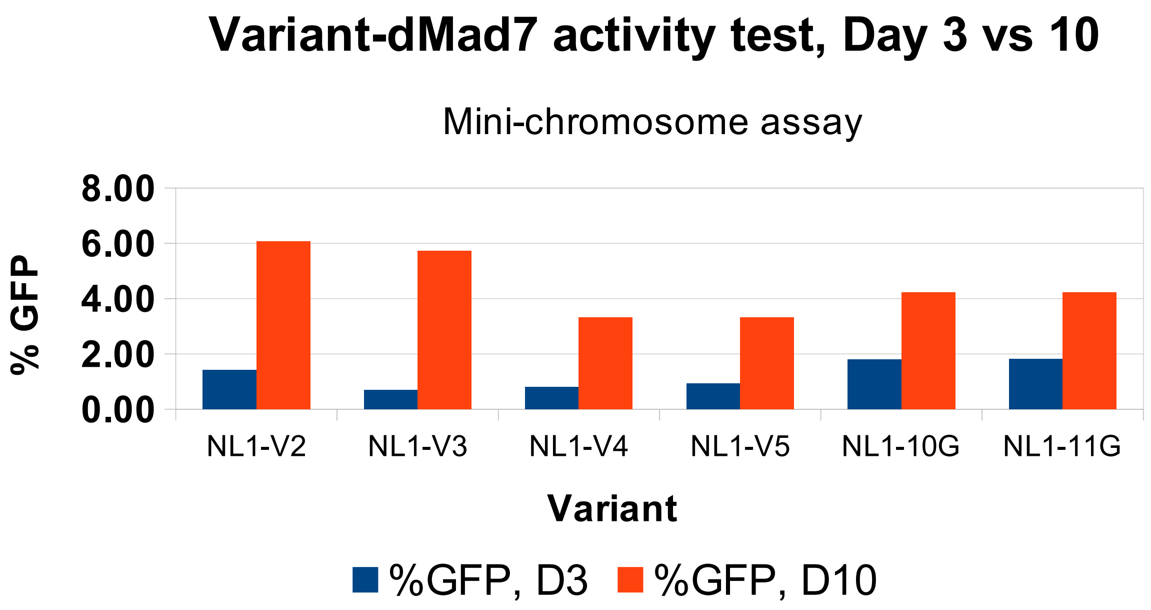


Stable-expression lines of variants V2, V3, V4, V5, 10G and 11G were transfected with the respective solo-expression plasmid, g10 and g6 expression plasmids, and an 3'intein-3'mCherry-2A-5'GFP donor. For each pool, GFP expression was checked in transfected cells (mCherry positive) at days 3 (D3) and 10 (D10) post-transfection. N=1.

**Figure S11:** Barcode skew

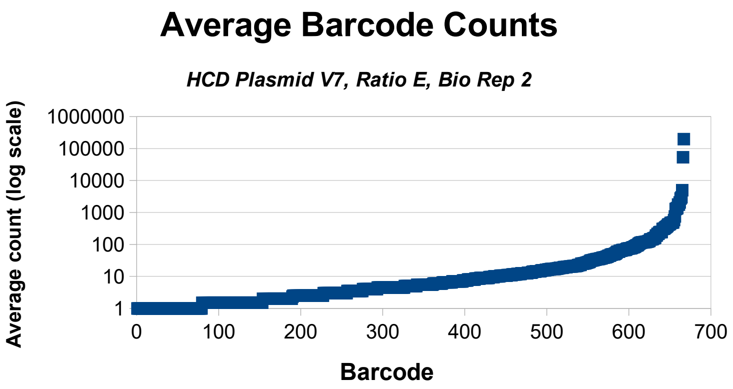


Plot of average barcode counts (log scale) for the transient native site A assay, High Cell Density V7, ratio E biological replicate 2. The barcodes were ordered by count in ascending order from left to right.

**Figure S12: Alteration of specificity assay**

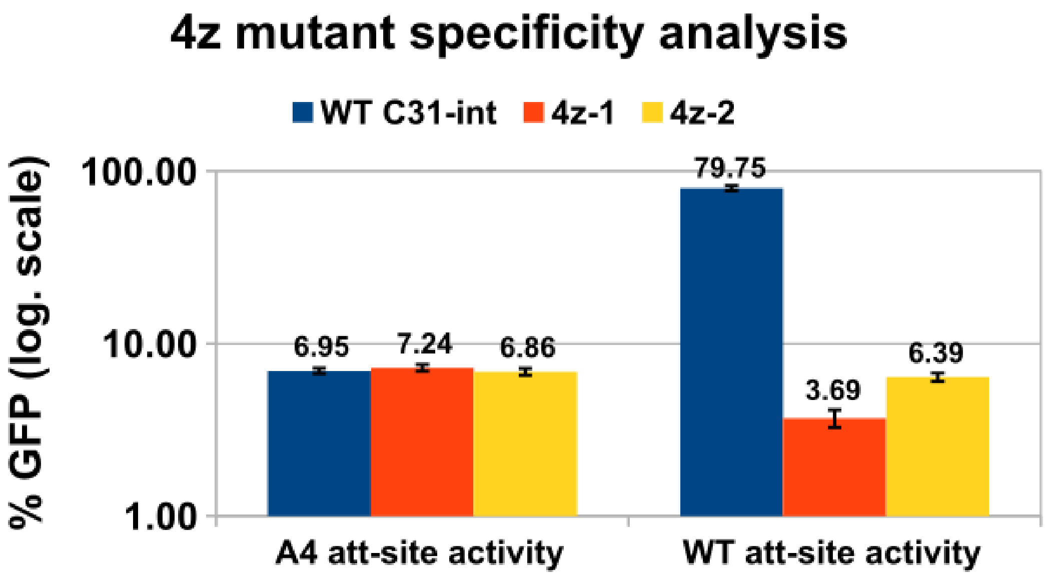


Activities of WT C31-int, variant 4z-1 and variant 4z-2 in the A4-intermediate and WT att-site inversion assays (left and right, respectively). Both variants were isolated via A4-intermediate screens. Logarithmic scale, N=3 biological replicates, error bars are standard error.
