## Supplementary Tables for "S-SELeCT: A Human-Evolved Serine Integrase System for Efficient Large-Cargo Genome Integration"

**Table S1**. Variant 7 mutation summary.

| Number | Mutation | AAs changed, deleted or inserted |
|---|---|---|
| 1 | EL -> FC at 162 | 2 |
| 2 | Y -> KK at 173 | 2 |
| 3 | L -> WV at 177 | 2 |
| 4 | 179 del SET | 3 |
| 5 | 230 del PFK | 3 |
| 6 | 233 ins LR | 2 |
| 7 | 274 del PA | 2 |
| 8 | VM -> GTF at 277 | 3 |
| 9 | EES -> PMW at 378 | 3 |
| 10 | R389K | 1 |

3.8% (23/605) AAs changed, deleted or inserted.

**Table S2**. Variant 12 mutation summary.

| Number | Mutation | AAs changed, deleted or inserted |
|---|---|---|
| 1 | V114A | 1 |
| 2 | I116A | 2 |
| 3 | GVFRQGNVMDLI -> RALRRGSIADPV at 122 | 9 |
| 4 | I136A | 1 |
| 5 | L139P | 1 |
| 6 | S142N | 1 |
| 7 | KES -> RGP at 144 | 3 |
| 8 | RE -> AF at 161 | 2 |
| 9 | A171R | 1 |
| 10 | Y -> KK at 173 | 2 |
| 11 | L -> WV at 177 | 2 |
| 12 | 179 del SET | 3 |
| 13 | INKLAH -> A at 195 | 5 |
| 14 | TT -> EPARSAA at 202 | 7 |
| 15 | T206A | 1 |
| 16 | H228R | 1 |
| 17 | 230 del PF | 2 |
| 18 | M251V | 1 |

| Number | Mutation | AAs changed, deleted or inserted |
|---|---|---|
| 19 | D254G | 1 |
| 20 | TR -> AW at 258 | 2 |
| 21 | PA -> TG at 274 | 2 |
| 22 | F289S | 1 |
| 23 | VIYKKK -> ATYRRG at 293 | 5 |
| 24 | D300G | 1 |
| 25 | 304 ins AG | 2 |
| 26 | 306 del KIE | 3 |
| 27 | Y -> GC at 310 | 2 |
| 28 | I312M | 1 |
| 29 | D315G | 1 |
| 30 | I317V | 1 |
| 31 | IE -> AG at 330 | 2 |
| 32 | WY -> RH at 335 | 2 |
| 33 | Q339R | 1 |
| 34 | D343G | 1 |
| 35 | G346D | 1 |
| 36 | K349A | 1 |
| 37 | 355 ins V | 1 |
| 38 | IL -> P at 357 | 2 |
| 39 | M361A | 1 |
| 40 | E367G | 1 |
| 41 | SK -> PE at 374 | 2 |
| 42 | EESIKDSYRCRR -> PIRVEGGCGCGE at 378 | 11 |
| 43 | K391G | 1 |

15.5% (94/605) AAs changed, deleted or inserted.

**Table S3**. Variant 29 mutation summary.

| Number | Mutation | AAs changed, deleted or inserted |
|---|---|---|
| 1 | E162F | 1 |
| 2 | A171R | 1 |
| 3 | Y -> KK at 173 | 2 |
| 4 | L177W | 1 |
| 5 | F -> LY at 231 | 2 |
| 6 | 274 del PA | 2 |
| 7 | VM -> GTIF at 277 | 4 |

| Number | Mutation | AAs changed, deleted or inserted |
|---|---|---|
| 8 | EES -> PMW at 378 | 3 |
| 9 | R389K | 1 |

2.8% (17/605) AAs changed, deleted or inserted.

**Table S4**. Variant 30 mutation summary.

| Number | Mutation | AAs changed, deleted or inserted |
|---|---|---|
| 1 | 161 del RELG | 4 |
| 2 | 165 ins F | 1 |
| 3 | A171R | 1 |
| 4 | 173 del YGF | 3 |
| 5 | L -> EW at 177 | 2 |
| 6 | 178 ins A | 1 |
| 7 | 230 del PFK | 3 |
| 8 | 233 ins LR | 2 |
| 9 | 274 del PA | 2 |
| 10 | VM -> GTF at 277 | 3 |
| 11 | EES -> PMW at 378 | 3 |
| 12 | R389K | 1 |

4.3% (26/605) AAs changed, deleted or inserted.

**Table S5**. Variant 32 mutation summary.

| Number | Mutation | AAs changed, deleted or inserted |
|---|---|---|
| 1 | V114A | 1 |
| 2 | VS -> AG at 117 | 2 |
| 3 | N128D | 1 |
| 4 | D131G | 1 |
| 5 | IH -> VR at 133 | 2 |
| 6 | M137V | 1 |
| 7 | LD -> PG at 139 | 2 |
| 8 | HK -> RR at 143 | 2 |
| 9 | S146P | 1 |
| 10 | REL -> GFC at 161 | 3 |
| 11 | 173 del YGF | 3 |
| 12 | L -> EW at 177 | 2 |
| 13 | 178 ins A | 1 |

| Number | Mutation | AAs changed, deleted or inserted |
|---|---|---|
| 14 | INK -> LGG at 195 | 3 |
| 15 | H200Y | 1 |
| 16 | T203A | 1 |
| 17 | LT -> PA at 205 | 2 |
| 18 | H228R | 1 |
| 19 | PFKP -> R at 230 | 4 |
| 20 | 274 del PA | 2 |
| 21 | VM -> GTIF at 277 | 4 |
| 22 | F289L | 1 |
| 23 | EVIYKKK -> RATHGEE at 292 | 7 |
| 24 | D300G | 1 |
| 25 | TTKIE -> AAEVG at 304 | 5 |
| 26 | I312A | 1 |
| 27 | D315G | 1 |
| 28 | I317T | 1 |
| 29 | 321 del PVE | 3 |
| 30 | DC -> AGPGH at 325 | 5 |
| 31 | IE -> AR at 330 | 2 |
| 32 | W335R | 1 |
| 33 | EL -> GP at 337 | 2 |
| 34 | L342S | 1 |
| 35 | 345 ins GG | 2 |
| 36 | 348 del GK | 2 |
| 37 | S352P | 1 |
| 38 | MDK -> VGG at 361 | 3 |
| 39 | CE -> RG at 366 | 2 |
| 40 | 370 ins IT | 2 |
| 41 | MTS -> P at 372 | 3 |
| 42 | EESIKD -> PMWVEG at 378 | 6 |
| 43 | 386 del RCR | 3 |
| 44 | 389 ins Y | 1 |
| 45 | 391 ins RE | 2 |

16.2% (98/605) AAs changed, deleted or inserted.

**Table S6**. Variant 36 mutation summary.

| Number | Mutation | AAs changed, deleted or inserted |
|---|---|---|
| 1 | REL -> GFD at 161 | 3 |
| 2 | A171R | 1 |
| 3 | Y -> KK at 173 | 2 |
| 4 | L -> WV at 177 | 2 |
| 5 | 179 del SET | 3 |
| 6 | H228R | 1 |
| 7 | 230 del PF | 2 |
| 8 | 274 del PA | 2 |
| 9 | 276 ins GTI | 3 |
| 10 | EES -> PMW at 378 | 3 |
| 11 | R389K | 1 |

3.8% (23/605) AAs changed, deleted or inserted.

**Table S7**. Variant 37 mutation summary.

| Number | Mutation | AAs changed, deleted or inserted |
|---|---|---|
| 1 | REL -> GFD at 161 | 3 |
| 2 | A171R | 1 |
| 3 | Y -> KK at 173 | 2 |
| 4 | L -> EWVA at 177 | 4 |
| 5 | 179 del SET | 3 |
| 6 | H228R | 1 |
| 7 | PF -> Y at 230 | 2 |
| 8 | 274 del PA | 2 |
| 9 | VM -> GTIF at 277 | 4 |
| 10 | EES -> PMW 378 | 3 |
| 11 | R389K | 1 |

4.3% (26/605) AAs changed, deleted or inserted.

**Table S8**. Variant 40 mutation summary.

| Number | Mutation | AAs changed, deleted or inserted |
|---|---|---|
| 1 | V114A | 1 |
| 2 | REL -> GFD at 161 | 3 |
| 3 | Y -> KK at 173 | 2 |
| 4 | 177 del LVS | 3 |
| 5 | 180 ins WVEV | 4 |

| Number | Mutation | AAs changed, deleted or inserted |
|---|---|---|
| 6 | H228R | 1 |
| 7 | F -> LY at 231 | 2 |
| 8 | 273 del DP | 2 |
| 9 | VM -> GTIF at 277 | 4 |
| 10 | D300G | 1 |
| 11 | D315N | 1 |
| 12 | I317T | 1 |
| 13 | I330V | 1 |
| 14 | L358P | 1 |
| 15 | EES -> PMW at 378 | 3 |
| 16 | R389K | 1 |

5.1% (31/605) AAs changed, deleted or inserted.

**Table S9**. Summary of total AAs changed, deleted or inserted.

| Variant | Total change count | Total change percent |
|---|---|---|
| V7 | 23 | 3.8% |
| V12 | 94 | 16.5% |
| V29 | 17 | 2.8% |
| V30 | 26 | 4.3% |
| V32 | 98 | 16.2% |
| V36 | 23 | 3.8% |
| V37 | 26 | 4.3% |
| V40 | 31 | 5.1% |

**Table S10**. Sequences targeted using Cas9 for chromatin accessibility DNA-segment knock-in assay

| **Locus code** | **Chromosome and band** | **Sequence of att-site candidate** | **SpCas9 target** | **PAM distance from att-site core [bp]** |
|---|---|---|---|---|
| A | 4p14 | GAGCATCCCCAACGAAGAGGACTTTCAGGTCTCCTTATTTGGGAAAACCC | CAAGAGCATCCCCAACGAAG | 8 |
| C | 9q34.3 | CTTATGCTCCAGGTGGATGCATGCACGAGTGTACCCAGCTGGGAGCTACC | CATGCACGAGTGTACCCAGC | 14 |
| D | 11p12b | GCTGTGCCTGAAGTATGGGGAGATTTAAGTCCTCATAGTTAACTTGGAAC | GAAGTTCCAAGTTAACTATG | 8 |
| E | 11q22.3 | TTATATGTCTAACCCAGAAAAGTGACTGTTTTTCTCATTTAAGGAAATAA | GTAAAATGGCCTGGTTCTAA | 147 |
