## Supplementary Materials and Methods for "S-SELeCT: A Human-Evolved Serine Integrase System for Efficient Large-Cargo Genome Integration"

#### Selection of Site A

We started the search for potential directed evolution targets by identifying all 50 bp human sequences that have a dyad symmetry (inverted semi-repeats) consistent with characterized serine integrases. Next, these sequences were aligned against the forward and reverse-complement phiC31 and Bxb1 attP sites. Candidates with a score in the top 75 were then manually screened for GC content between 20-60%, uniqueness, overlap with known genes, proximity to DNAseI hypersensitivity sites (HSS), and information on the flanking genes. A set of 17 active-integrase attP sites was used to determine the GC content boundaries: phiC31, phiTB1, R4, Bxb1, TP901, TG1, Int2, Int3, Int4, Int5, Int7, Int8, Int9, Int10, Int11, Int12, and Int13 (1–7). BLAT was used to check for uniqueness (8). To investigate gene overlap, DNaseI HSS distance, and neighboring gene information, we used the UCSC Genome Browser (GRCh38/hg38) (9). To assess if the flanking genes were known to be potential direct oncogenes, we performed PubMed searches. For prediction of expression, we consulted GTEx RNA-Seq expression data of the 5′ and 3′ genetic neighbors (10). The DNAseI HSS data that we used was from the ENCODE project (11). To further test the openness of chromatin at the candidate sites, we confirmed that Cas9 was able to mediate knock-in of a 4 kb insert (CMV-puro-TK-GFP) at sites A, C, D and E in HEK293 cells (NHEJ, pool-based PCR analysis, Table S10 and data not shown).

#### Intronization

To intronize sequences, we first performed codon optimization using the GeneArt GeneOptimizer tool, with desired restriction sites and strong potential exon splice junctions excluded (CAG/GT, AAG/GT) (12). Next, we created a custom algorithm to introduce exon splice junctions at desired intervals via silent codon changes. We weighted potential splice junctions with guidance from the Ma et al. sequence logos (i.e. high scoring for 5′ C/A-A-G, weak preference for 3′ G-T-T) (12). Our exons mostly fall in the 200-300 nt range. Finally, we inserted introns at the introduced exon splice junctions. Introns were obtained from endogenous human genes including: ACTB, GAPDH, PGK1, PPIA, RPL13A, RPLP0, TTN, and YWHAZ. They were either naturally less than 201 nt in size, or we truncated them to meet this requirement (by fusing together the first and last 100 nt of the intron). In certain instances, we modified introns to start with the dominant "GTAAG" motif identified by Ma et al. (12). We have annotated all exons and introns in our deposited sequences.

#### General plasmid cloning

All enzymes used for cloning PCRs, restriction-digests, ligations and Golden Gate Assembly (GGA) were purchased from NEB. We used the GeneArt GeneOptimizer® tool for codon optimization (13). DNA segments were ordered from GeneArt when they contained codon optimizations produced by their tool. In all other cases, dsDNA segments and oligos used for cloning were purchased from IDT. PCR was performed with Q5 Hot Start High-Fidelity 2X Master Mix (NEB M0494L). DNA segments were gel- or column-purified using Zymo Research DNA Clean & Concentrator kits and reagents (e.g. D4004), then quantified using the Qubit 1X dsDNA Broad Range kit (Thermo Q33266) or UV-absorption (BioTek Synergy HTX). For elution from columns, Qiagen Buffer EB (Qiagen 19086) was used (heated to 95°C for segments larger than 2 kb).

When performing cloning via ligation of purified restriction-enzyme-digested segments, we combined 5 femtomoles (fmol) of plasmid backbone with 15 fmol of each insert in a 5 uL reaction. These mixes contained 0.5 uL 10X T4 DNA Ligase Buffer, 0.5 uL 10 mM ATP (NEB P0756S) and 0.5 uL T4 DNA Ligase (400 U/uL; NEB M0202L). When only one insert was present, we thermocycled the ligation reaction as follows: 15 cycles of 10°C for 30 sec, 30°C for 30 sec; 1 cycle of 65°C for 12 min, 4°C hold. For higher insert counts, we increased the cycling: 30 cycles for two inserts, 45 cycles for three inserts, etc.

For GGA, we primarily used the NEB BsaI-HFv2 and BsmBI-v2 Golden Gate enzyme mixes (NEB E1601L and E1602L, respectively). For 1-3 insert GGA, we combined 5 fmol of plasmid backbone with 15 fmol of each insert in a 5 uL reaction. The mixes contained 0.5 uL 10X T4 DNA Ligase Buffer and 0.25 uL of the BsaI or BsmBI GGA enzyme mix. When only one insert was present, we thermocycled BsaI GGA reactions as follows: 15 cycles of 37°C for 30 sec, 16°C for 30 sec; 1 cycle of 65°C for 12 min, 4°C hold. BsmBI GGA single-insert reactions were cycled between 42°C and 16°C, i.e.: 15 cycles of 42°C for 30 sec, 16°C for 30 sec; 1 cycle of 65°C for 12 min, 4°C hold. For higher insert counts, we increased the cycling: 30 cycles for two inserts, 45 cycles for three inserts.

To perform GGA involving 4 or more inserts, we increased the volumes and DNA amounts as needed, and switched to an equal molecular ratio (i.e. all backbone and inserts present at the same copy number). Final DNA concentrations ranged between 1-6 nM, and cycling conditions were increased up to the manufacturer's recommended maximum (30 cycles of 37°C/42°C for 5 min, 16°C for 5 min); reactions were then heat-killed at 65°C for 12 minutes and held at 4°C. When a different type IIS restriction enzyme was needed, we used the NEBridge Ligase Master Mix (NEB M1100L) and followed the manufacturer's guidance.

After ligation or GGA, we combined 1 uL of the reaction with 20 uL of chemically-competent 10-beta E. coli cells (NEB C3019H) in a 300 uL thin-walled PCR tube, and then incubated the solution at 4°C for 30 minutes. Transformation was performed by heat-shocking the cells for 30 seconds in a thermocycler heat-block set to 42°C. The cells were immediately chilled at 4°C for 5 minutes, then transferred into 400 uL of NEB 10-beta/Stable Outgrowth Medium (NEB B9035S). The cells were grown for 1 hour at 37°C with 250-300 RPM shaking, then spread onto LB Agar plates with the needed antibiotic, and incubated for 12-18 hours at 37°C.

**Plasmid DNA amplification, purification and sequencing**

For all scales of plasmid amplification, we started by seeding a single colony into 2 mL of Supreme Media (SM) or Terrific Broth (TB; Teknova T7660) supplemented with the needed antibiotic: 50 ug/mL kanamycin (Fisher AAJ67354AD), 100 ug/mL carbenicillin (Teknova C2199), or 34 ug/mL chloramphenicol (Fisher AAJ67273AB). The growth was incubated at 37°C for 16-24 hours with shaking at 250-300 RPM. 500 mL SM was prepared as follows: 472.51 mL NERL High Purity Water (Fisher 23-249-590), 4.13 mL 50% (w/w) Glucose (Teknova G9005), 18.11 g Bacto CD Supreme (Fisher A4973701), 5 mL 99.5% Glycerol (Fisher AC327255000), 0.25 g L-Leucine (MP Biomedicals 02102158.3). The medium was then filter-sterilized (Corning 431097) and stored at 4°C.

Mini-prep scale plasmid purification was performed using the QIAprep Spin Miniprep Kit (Qiagen 27106) and protocol, however with doubled volumes. I.e. 2 mL cell growths were pelleted, resuspended in 500 uL Buffer P1, lysed with 500 uL Buffer P2 for 3-5 minutes, then neutralized with 700 uL Buffer N3. The lysate was centrifuged for 10 minutes at 20,000 RCF, then decanted into a separate 2 mL tube. Clarified lysate was passed through a single QIAprep column via centrifugation (18,000 RCF for 1 min) or vacuum pressure (300 mbar), and the column was then washed with 0.8-1 mL Buffer PE. Drying was performed via centrifugation (18,000 RCF for 1 min) and incubation at 50°C for 5 minutes, both in fresh collection tubes. For elution, we used 30-60 uL of Buffer EB.

For midi-prep scale purification, we seeded 300 uL of the 2 mL growth into 140 mL of antibiotic-supplemented SM or TB in a sterile 250 mL PETG baffled Erlenmeyer flask (Thermo 10-530). If using carbenicillin, before seeding we first washed the cells twice with fresh growth medium (because the resistance enzyme is secreted/present outside the cells). Flasks were then incubated at 37°C for 16-24 hours with 250-300 RPM shaking. Growths were aliquoted into pre-weighed 50 mL tubes, then centrifuged at 3000 RCF for 15 minutes at 4°C. After discarding supernatants, tubes were weighed again to determine pellet masses. We next used the NucleoBond Xtra Midi EF and Finisher Midi kits (Machery-Nagel 740420.50 and 740439.50) according to the manufacturer's protocol, however with modified wash and elution steps. 10.67 mL of each buffer (RES-EF, LYS-EF, NEU-EF) was used per 1 gram of cell pellet. Both wash steps were repeated an additional time (i.e. two 35 mL ENDO-EF and two 15 mL WASH-EF washes in total). Qiagen Buffer EB heated to 50°C was used to elute from finisher columns, and concentrations were determined via UV-vis absorption (BioTek Synergy HTX). Frequently used plasmids were aliquoted and stored at -20°C in a non-cycling freezer (i.e. not frost-free).

All plasmids were sequenced with Oxford Nanopore devices, and custom scripts were used to analyze the results (alignment to expected sequence, identification of mismatches, coverage level, etc.). Examples of products used include the Rapid Barcoding Kit 24 V14 (Oxford Nanopore Tech. SQK-RBK114.24), R10.4.1 flongle flow cells (FLO-FLG114), and R10.4.1 MinION flow cells (FLO-MIN114).

**General library cloning**

GGA was used to create all libraries. We most often created a 30 uL mix with 90 or 180 fmol of each segment present, i.e. 3 or 6 nM final concentrations. The solutions contained 3 uL 10X T4 DNA Ligase Buffer and 1.5 uL of the BsaI-HFv2 or BsmBI-v2 Golden Gate enzyme mix. We used a thermocycler to incubate BsaI GGA reactions as follows: 30 cycles of 37°C for 5 min, 16°C for 5 min; 1 cycle of 65°C for 12 min, 4°C hold. BsmBI GGA reactions were incubated at 42°C instead of 37°C: 30 cycles of 42°C for 5 min, 16°C for 5 min; 1 cycle of 65°C for 12 min, 4°C hold. After GGA, the reactions were purified and concentrated via column-purification using the Zymo DNA Clean & Concentrator kit (Zymo D4004). We followed the manufacturer's protocol, however to elute the DNA we used 8 uL of Buffer EB heated to 95°C (Qiagen 19086).

Next, 1-3 uL of the purified GGA was combined with 25 uL of 10-beta E. coli (NEB C3020K) in a 0.1 cm gap cuvette (Bio-Rad 1652083), and the mix was electroporated using a BTX Electro Cell Manipulator 600 at 1.7 kV with 186 Ohm resistance. If successful (i.e. no spark, 7-8 msec discharge time), the cells were then immediately transferred into 1 mL of NEB 10-beta/Stable Outgrowth Medium (NEB B9035S), and incubated for 1 hour at 37°C with 250-300 RPM shaking. Serial dilutions of the outgrowth were plated for library size or coverage estimates. The remainder of the growth (typically 925-950 uL; specific amount measured) was then seeded into 140 mL of antibiotic-supplemented SM or TB in a sterile 250 mL PETG baffled Erlenmeyer flask (Thermo 10-530). The cells were incubated at 37°C for

16-24 hours with shaking at 250-300 RPM. If the library size/coverage was adequate, we then purified the plasmid pool using our described midi-prep protocol.

**Inversion-assay plasmid assembly**

Three-exon GFP recombination-reporter plasmids were cloned using Golden Gate Assembly (GGA) of synthesized DNA segments and PCR products (14). They consist of two expression cassettes oriented in a divergent manner, flanked by UMS terminator sequences (15). The first is driven by a CMV promoter, contains the three-exon GFP segment, and is terminated by the Bovine Growth Hormone (BGH) 3′ UTR/polyA. The two introns in GFP are derived from Li et al. (16) and the second human Hemoglobin Subunit Beta intron (HBB; NCBI Gene 3043). The second expression cassette consists of the human EF1-alpha promoter, a 3′ intein-3′ mCherry2 protein fusion, WPRE (Woodchuck Hepatitis Virus Posttranscriptional Regulatory Element), and an SV40 3′ UTR/polyA (17–19). We have deposited the sequences of a representative plasmid sequence (pKSP3-dAf-8) and the corresponding recombination product (pKSP3-dAf-8-flipped) at Zenodo.

**Mini-chromosome assay reporter plasmid cloning**

Mini-chromosome assay reporter plasmids were cloned using GGA of synthesized DNA segments and PCR products. They contain a split-GFP integration reporter, a polycistronic trEBNA-1 puroR expression cassette, and the EBV oriP replication origin (20). The expression cassette is driven by the human EF1-alpha promoter, contains a WPRE, and is terminated by the BGH 3′UTR/polyA. trEBNA-1 is a truncated version of the EBNA1 protein (20), and it is separated from the puromycin-resistance protein via P2A-mediated ribosomal skipping (21). The integration reporters consist of an attP-3′ intron-3′GFP-BGH 3′UTR/polyA segment that is flanked by genomic Site A sequence. The attP is either from WT phiC31 or the human Site A sequence. The 3′ intron is derived from the synthetic intron acceptor described by Li et al. (16), and 3′GFP is a C-terminal segment from EGFP4m (mut4EGFP) (22). In all plasmids, upstream and downstream Site A genomic sequences have been cloned 5′ and 3′ of the reporter, respectively. In the MAX and SAR1 vectors, 5 kb and 2.5 kb genomic sequence has been cloned in each region (10 and 5 kb total in each plasmid), respectively. A 1.2 kb chicken heat-shock 4 (cHS4) insulator (23) separates the reporter and trEBNA-1 puroR expression cassette. Sequences of the following reporter plasmids have been deposited at Zenodo: pKEO-SAR1-WT-attP, pKEO-SAR1, pKEO-SAR-MAX-WT-attP, pKEO-SAR-MAX.

**Split-GFP assay donor plasmid assembly**

Donor plasmids used in both chromosomal and mini-chromosome split-GFP assays were cloned using GGA of synthesized DNA segments and PCR products. They contain a single CMV-driven polycistronic cassette that is terminated by an SV40 3′UTR/polyA. The first protein expressed is a 3′ intein-3′ mCherry2 protein fusion (17, 18). The next protein is separated via a P2A peptide, and consists of an N-terminal EGFP4m exon without a stop codon. An intron splice donor derived from Li et al. (16) borders the EGFP4m exon, and the following elements are downstream of it: (i) either the wildtype or fully mutated Site A attB, (ii) an intron splice acceptor derived from HBB, (iii) the sLTSV(-) hammerhead ribozyme (HHR) (24, 25). Both plasmid sequences (pDiG-SP3-WT-attB, pDiG-SP3-Af8) have been deposited at Zenodo.

**Native Site A donor plasmid cloning and barcoding**

The native Site A donor precursor-plasmid was cloned using GGA of synthesized DNA segments and PCR products. It contains a single CAG-driven heavily-intronized polycistronic cassette with a WPRE and BGH 3'UTR/polyA terminator. Three proteins are co-expressed and separated via P2A peptides: EGFP4m, a 3' intein-3' mCherry2 fusion, and a puromycin-resistance enzyme (17, 18, 22). The cassette is flanked by a single full-length cHS4 insulator on each side (23). Upstream of the first insulator, the plasmid also contains a SapI Golden Gate cloning platform, and the fully mutated Site A attB sequence. We have deposited sequences of the precursor (pFAD-HBH) and barcoded (degenerate) donor (pFAD-HBH-K18) at Zenodo.

To create a barcoded plasmid pool, we synthesized a degenerate oligonucleotide mix, extended it with Q5 polymerase to create dsDNA, then performed Golden Gate Assembly with SapI. The degenerate oligonucleotide mix was synthesized as an IDT Ultramer, and had the following sequence: 5'-gaaaaggagaagggaaagagggaggttgttGCTCTTCtAGAa KKKKKKaagaaKKKKKKaggaKKKKKKaGAGtGAA GAGCttgttgTGAGATCTTTTGATCTCAcaacaagctcttcactc-3'. To produce double-stranded DNA (dsDNA), we first created the following 100 uL mix: 30 uL UltraPure H2O (Thermo), 50 uL Q5 HotStart HF 2X Master Mix (NEB M0494L), 20 uL 10 uM oligo. There is no second primer because the oligo forms a 3' hairpin (GAGtGAAGAGCttgttgTGAGATC TTTT GATCTCAcaacaagctcttcactc). Next, the mix was thermocycled as follows: 98°C for 30 sec; 10 cycles of 98°C for 7 sec, 65°C for 10 sec (-0.5°C/cycle), 72°C for 4 sec; 25 cycles of 98°C for 7 sec, 60°C for 10 sec, 72°C for 4 sec; 72°C for 2 min, 4°C hold. The ~94 basepair (bp) product and 10.1 kbp SapI-digested plasmid backbone were then gel-purified, and used in the following 30 uL Golden Gate Assembly (GGA) mix: 11.36 uL UltraPure H2O, 10 uL 3X NEBridge Ligase Master Mix (NEB M1100S), 3.64 uL backbone (49.4 fmol / uL), 1 uL dsDNA barcode pool (180 fmol / uL), 4 uL SapI (NEB R0569S). The mix was thermocycled as follows: 37°C for 5 min; 30 cycles of 16°C for 1 min, 37°C for 1 min; 4°C hold. Next, we column-purified the mix according to the manufacturer's protocol (Zymo D4004), however for elution we used 8 uL of Qiagen Buffer EB heated to 95°C. We electroporated 3 uL of the purified GGA into 25 uL of electrocompetent cells (NEB C3020K) using 0.1 cm gap cuvettes (Bio-Rad 1652083) and a BTX Electro Cell Manipulator 600 (1.7 kV, 186 Ohm), then followed the remainder of our standard plasmid library production protocol. Based on colony-forming-unit (CFU) counts of the post-electroporation 1 hour outgrowths, our barcoded-plasmid pool diversities ranged from 232-259 e6.

**Solo-integrase stable-expression plasmid assembly**

The precursor plasmid used for non-fused expression of WT C31-int and variants was cloned using GGA of synthesized DNA segments and PCR products. It contains two divergent promoterless expression cassettes centered on a Bxb1 attB site. The first incomplete cassette contains the coding sequence for a blasticidin-resistance protein, and it is terminated by an SV40 3'UTR/polyA. The second partial-cassette has a BsaI GGA platform flanked by WT C31-int coding sequence with a C-terminal mouse TCOF NLS. Downstream of this, the following elements are present: P2A peptide, 5' mCherry2-5' intein fusion protein, BGH 3'UTR/polyA. In addition, two UMS terminators are present in the plasmid, one on each flank of the divergent partial-cassette segment. To clone complete integrase sequences or variant libraries, BsaI GGA was performed using our standard conditions (for plasmids and libraries, respectively). We have deposited the precursor (pHKi-CL-357-60mT), WT C31-int (pHKi-WTC31-60mT), V7 (pHKi-V7-60mT), and V12 (pHKi-V12-60mT) plasmid sequences at Zenodo.

### Transient solo-integrase expression plasmid cloning

For transient-expression of variants in a non-fused form, a precursor plasmid was cloned using GGA involving synthesized DNA segments and PCR products. Most plasmid backbone segments were obtained from pCMV-SPORT-βgal (Thermo 10586014) via PCR. The precursor vector has a single expression cassette driven by a CMV promoter, and it contains the following elements: BsaI GGA platform with flanking WT C31-int coding sequence, C-terminal human TCOF NLS, 3′ small SV40T intron (from pCMV-SPORT-βgal), SV40 3′UTR/polyA. To clone variants, a central segment encoding the desired mutations was generated via PCR or gene synthesis, then it was cloned into the precursor plasmid backbone (BsaI digested and gel-purified) using our standard BsaI GGA conditions. We have deposited the precursor vector (pDCMV-CL-357-hT60), V7, V29, V30, V32, V36, V37 and V40 solo-expression plasmid sequences at Zenodo (pDCMV-V7-hT60, pDCMV-V29-hT60, pDCMV-V30-hT60, pDCMV-V32-hT60, pDCMV-V36-hT60, pDCMV-V37-hT60, and pDCMV-V40-hT60, respectively).

### Fused-integrase stable-expression plasmid assembly

The precursor plasmid used for expression of WT C31-int and variants fused to dMad7 was cloned using GGA of synthesized DNA segments and PCR products. It contains two divergent promoterless expression cassettes centered on a Bxb1 attB site. The first incomplete cassette contains the coding sequence for a blasticidin-resistance protein, and it is terminated by a BGH 3′UTR/polyA. The second partial-cassette has a BsaI GGA platform flanked by intronized WT C31-int coding sequence. Downstream of this is the NL1 or NL2 NLS-linker that is used to fuse integrase to intronized dMad7. The NL1 sequence consists of the mouse TCOF NLS and a $Gly_4$-Ser-Gly-Gly linker, whereas NL2 is a fusion of the human Nucleoplasmin NLS and an $Gly_4$-Ser-$Gly_4$-Ser-Gly-Gly-Gly linker. Two SV40 NLS tags are present at the C-terminus of dMad7, and the following elements are located immediately downstream: T2A peptide, intronized 5′ mCherry2-5′ intein fusion protein, WPRE, SV40 3′UTR/polyA. In addition, two UMS terminators are present in the plasmid, one on each flank of the divergent partial-cassette segment. To clone complete integrase sequences, BsaI GGA was performed using our standard conditions. We have deposited the NL1-precursor (pLKi-CL357-NL1-dM7), NL2-precursor (pLKi-CL357-NL2-dM7), and V30-NL2 (pLKi-V30-NL2-dM7) plasmid sequences at Zenodo.

### Assembly of stable-expression alternative-splicing solo and fused integrase plasmids

Precursors of vectors used to simultaneously stably-express solo and dMad7-fused integrases were cloned via GGA of synthesized DNA segments and PCR products. They contain two divergent promoterless expression cassettes centered on a Bxb1 attB site. The first incomplete cassette contains the coding sequence for a blasticidin-resistance protein, and it is terminated by a BGH 3′UTR/polyA. The second partial-cassettes contain the following elements: a BsaI GGA platform flanked by intronized WT C31-int coding sequence, splice donor, splice-1 acceptor, human TCOF NLS, BGH 3′UTR/polyA, splice-2 acceptor, NL1 or NL2 NLS-linker, intronized dMad7, two SV40 NLS tags, T2A peptide, intronized 5′ mCherry2-5′ intein fusion protein, WPRE, SV40 3′UTR/polyA. In addition, two UMS terminators are present in the plasmid, one on each flank of the divergent partial-cassette segment. To clone complete integrase sequences or variant libraries, BsaI GGA was performed using our standard conditions (for plasmids and libraries, respectively). We have deposited the NL1-precursor (pLKa4-

NL1-dM7-plat), NL2-precursor (pLKa4-NL2-dM7-plat), and V12-NL1 (pLKa4-V12-NL1-dM7) plasmid sequences at Zenodo.

**gRNA identification and expression plasmid cloning**

Potential guide RNAs were identified using the CRISPOR tool with the YTTV PAM Mad7 setting (26). The precursor of the vectors used to transiently express guide RNAs was assembled via GGA of synthesized DNA and PCR products. There is a single cassette in the plasmid that contains the human U6 promoter, 'TAATTTCTACTCTTGTAGAT' crRNA scaffold (27), and a BsmBI GGA platform. To clone gRNA-expression plasmids, BsmBI GGA was performed. The g6 and g10 expression vectors were made by cloning the 'TGTGGGAGTTCAGTTCAACAACT' and 'AGCCGCTACCCCGACCACATGAA' spacers, respectively. We deposited the precursor (pKU6-M7-plat), g6 (pKU6-M7-SA-6) and g10 (pKU6-M7-SA-10) plasmid sequences at Zenodo.

**HEK293 cell culture**

HEK293 cells were maintained in complete DMEM medium consisting of high-glucose DMEM (Fisher 11-960-077) supplemented with 4.5% fetal bovine serum (FBS; Corning 35-075-CV), 2 mM GlutaMAX (Fisher 35-050-061), 1 mM sodium pyruvate (Fisher 11-360-070), 50 mM HEPES (Fisher 15-630-080), 100 ug/mL Normocin (InvivoGen ant-nr-1), 50 units/mL penicillin, and 50 ug/mL streptomycin (Fisher 15-140-148). CellBIND multi-well plates and flasks were used at all times to improve cell attachment (e.g. Corning 3337, 3335, 3290, 3291, etc.). Cells were cultured at 37°C in a humidified incubator with 5% $CO_2$ (Fisher Scientific, Model 3530). To improve survival after FACS isolation, FBS was increased to 10% final for 1 week, then returned to 4.5%. Cells that contained the H11 landing-pad were sometimes grown with 200 ug/mL hygromycin (InvivoGen ant-hg-1) present. Libraries were maintained via selection with 20 ug/mL blasticidin (InvivoGen ant-bl-1). Cultures of cell lines with the SAR1 or SARMAX mini-chromosome were additionally supplemented with 1-2 ug/mL puromycin (InvivoGen ant-pr-1). When both hygromycin and puromycin were present, we also added 2.5 mM oxaloacetic acid (Fisher AA1578909) to improve cell growth and health. Cells were passaged at 70-90% confluency using TrypLE Express (Fisher 12-605-028) with 1:20-1:2 split densities.

**Transfection**

The day before transfection, cells were seeded with the aim of reaching 60-80% confluency in 18-24 hours. In experiments involving transient expression, plain cell culture medium without added drugs was used (high-glucose DMEM, 4.5% FBS, 2 mM GlutaMAX, 1 mM sodium pyruvate, 50 mM HEPES), and the final volume per well of a CellBIND 24 well-plate (24ww; 2 $cm^2$) was 0.5 mL. When expression occurred from the H11 landing-pad, cells were seeded in plain cell culture medium with 1 ug/mL doxycycline (Sigma D3072-1ML). Plain HEK293 and H11-landing-pad cells were seeded at densities of 175-225k live cells per 24ww. Cells with the SAR1 or SARMAX mini-chromosome were seeded at densities in the 300-350k range per 24ww. Cell counts were performed with a Countess II (Thermo) in the presence of 0.2% trypan blue live/dead stain.

The poly beta-amino ester transfection reagent Xfect (Takara 631318) was used for all plasmid delivery, at a 0.3 uL per 1 ug ratio. At minimum, 3.33 ug of plasmid DNA was mixed with 1 uL Xfect in a final volume of 55.4 uL. The solution was vortexed well immediately after addition of Xfect, then complexes were allowed to form for 10-15 minutes. Before pipetting, the sample was briefly vortexed, then 33.3 uL (2 ug DNA) was delivered into each 24 ww. These plasmid amounts, cell densities and volumes were proportionally scaled up or down when transfecting at different scales (e.g. everything increased 5X when working in 6 well-plates, which have 10 $cm^2$ growth area).

**Creation of split-GFP WT-attP reporter line**

To introduce split-GFP reporter sequences at both Site A loci, we took a four-step approach where full-cassette knock-in was followed by excision of the promoter and 5′ segment of GFP. First, an attL-CMV-5′ GFP-attR-3′ GFP-BGHpA amplicon from a pool was knocked-in to a Site A locus using SpCas9-stimulated NHEJ. PCR from a plasmid template was used to create the insert pool with the following primers: 5′-g*t*tgtt NNNNNN ggTAGGGATAACAGGGTAAT ggctacgcccccaac tgagagaac*t*c-3′, 5′-c*a*acaa NNNNNN AAGCATGCCTGCTATTGTCTTC CCAATC-3′, where '*' is used to indicate a phosphorothioate linkage. The sequence of the PCR amplicon (EXLP1_amplicon) has been deposited at Zenodo. SpCas9 and the guide RNA were delivered in RNP form, and cleavage was targeted at the following spacer: CAAGAGCATCCCCAACGAAG (AGG PAM). After two weeks of outgrowth, GFP-positive cells were sorted into a pool. Second step: after 1 week of growth, these cells were then transfected with pCS-kRI, which co-expresses the phiC31 RDF and integrase (28). Two weeks after transfection, single GFP-negative cells were sorted, outgrown and screened for a desirable attP-3′ GFP-BGHpA knock-in via genotyping.

Our third and fourth steps were to repeat this process at the second locus with a slightly different insert, and an alternate spacer. The sequence of the PCR amplicon (EXLP2_amplicon) has been deposited at Zenodo, and the primers used to generate it were: 5′-g*t*tgtt NNNNNN ctacgcccccaactgagagaac*t*c-3′, 5′-c*a*acaa NNNNNN AAGCATGCCTG CTATTGTCTTCCCAATC-3′. The spacer used was: GACTTTCAGGTCTCCTTATT (TGG PAM). The sequences of the first and second locus in the clone that we chose have been deposited at Zenodo (SG51d6__first_locus, SG51d6__second_locus).

**WT-attP Site A integration efficiency assay**

When using transiently-expressed C31-int vectors to measure integration efficiency in our Site A WT-attP split-GFP assay, we used a 1:3 integrase:donor plasmid ratio. I.e. in a 24 ww, Xfect transfection was used to deliver 0.5 ug and 1.5 ug of the integrase and donor vectors, respectively. When solo C31-int was expressed from the H11 locus (e.g. in NLS screens), we transfected 2 ug of the donor plasmid per 24 ww, and 10 ug per 6 ww. For stably-expressed C31-int-dCas fusions (Sp dCas9 or dMad7), gRNA-expression, solo-integrase expression, and donor plasmids were present at 6%, 19% and 75% of the DNA mix by mass, respectively. Unless otherwise indicated, percentages of cells positive for both mCherry and GFP were measured 72 hours after transfection via fluorescent flow-cytometry (BD FACSMelody).

**Inversion assay**

The three-exon GFP inversion assay was only conducted in cells that expressed a solo or fused integrase from the H11 locus. We transfected 2 ug of the reporter plasmid per 24 ww, and 10 ug per 6 ww. Unless otherwise indicated, percentages of cells positive for both mCherry and GFP were measured 72 hours after transfection via fluorescent flow-cytometry (BD FACSMelody).

**Mini-chromosome establishment and assay**

To create pools of HEK293 H11 LP cells with a mini-chromosome reporter vector (e.g. SAR1, SARMAX), we first used Xfect to transfect 2 ug of the respective plasmid per 24 ww. Three days after transfection, we then split the cells at a 1:2-1:4 ratio, and cultured them in medium supplemented with 1-2 ug/mL puromycin, 200 ug/mL hygromycin, and 2.5 mM oxaloacetic acid for a minimum of two weeks. After this period, splits were performed at a 1:10 ratio. These supplements were only removed when performing integration efficiency tests.

To conduct the reporter assay in lines that express integrase fusions from the H11 locus, we used the same 6%-19%-75% DNA mix described for the WT-attP Site A efficiency tests. When using alternative-splicing to express both solo and fused integrases from H11, the gRNA-expression and donor plasmids constituted 6% and 94% of the DNA mix delivered per well, respectively. When expressing variant-dMad7 fusions transiently, we tested a variety of plasmid ratios. For two-plasmid experiments, the alternative-splicing integrase-dMad7 to donor mass ratios were as follows: E=1:3, F=1:2, G=1:1, H=2:1, i=3:1. In three-plasmid experiments, we tested the following fused : solo : donor ratios: J= 1:1:1, K=1:1:2, L=2:1:2, M=2:1:3, N=2:1:1. Unless otherwise indicated, percentages of cells positive for both mCherry and GFP were measured 72 hours after transfection via fluorescent flow-cytometry (BD FACSMelody).

**Native Site A assay**

In cells that express integrase fusions from the H11 locus, to perform the native Site A assay, we again used the 6%-19%-75% DNA mix for gRNA-expression (equal mix of g6 and g10), solo integrase-expression and donor plasmids, respectively. When alternative-splicing was used to express solo and fused integrase from H11, the gRNA-expression and donor plasmids were 6% and 94% of the DNA mix delivered per well, respectively. For transient expression tests, we used the same ratios and labels established for the mini-chromosome assay.

Unless otherwise indicated, cells positive for both mCherry and GFP were sorted 72 hours after transfection. In stable-expression experiments, 61.2 – 90k cells were sorted per 24 ww, and they were then outgrown for a week before being pelleted. In transient-expression tests, 5k cells were sorted per 24 ww, outgrown for a week in normal sort medium, then cultured in 1 ug/mL puromycin-supplemented medium for 1-2 weeks until the well was 80-100% confluent, after which they were pelleted.

Genomic DNA (gDNA) was extracted from frozen cell pellets using the Quick-DNA Microprep kit (Zymo Research D3021) according to the manufacturer's instructions, however we eluted using Qiagen Buffer EB. DNA concentrations were measured using the Qubit 1X dsDNA Broad Range (BR) kit. Next, we performed the primary 1199 bp PCR with primers 5'-CGCCTTCTATCGCCTTCTT*G*A-3' and 5'-CCTCACCTCACATGGCTC*T*G-3', where * indicates phosphorothioate linkages. The solution contained 50 ng gDNA, 1.25 uL 10 uM forward primer, 1.25 uL 10 uM reverse primer, 12.5 uL Q5 HotStart 2X Master Mix, and was brought to 25 uL with UltraPure H2O (Fisher 10-977-015). The thermocycling conditions were as follows: 1 cycle of 98°C for 1 min; 10 cycles of 98°C for 14 sec, 67°C for 20 sec (decrease 0.5°C/cycle), 72°C for 51 sec; 25 cycles of 98°C for 7 sec, 62°C for 10 sec, 72°C for 51 sec; 1

cycle of 72°C for 5 min. If the desired 1199 bp band was visible in an agarose gel run, we then performed nested 1039 bp PCR with primers 5′-ACTTTTCGGGGAAATGTG*C*G-3′ and 5′-GCATTCTTGCAGCAATCTC*G*G-3′. The solution contained 2 uL 1:100 diluted 1199 bp PCR, 2.5 uL 10 uM forward primer, 2.5 uL 10 uM reverse primer, 25 uL Q5 HotStart 2X Master Mix, and 18 uL UltraPure $H_2O$. The cycling conditions were as follows: 1 cycle of 98°C for 1 min; 10 cycles of 98°C for 14 sec, 66°C for 20 sec (decrease 0.5°C/cycle), 72°C for 42 sec; 25 cycles of 98°C for 7 sec, 61°C for 10 sec, 72°C for 42 sec; 1 cycle of 72°C for 5 min. If the desired 1039 bp band was visible in an agarose gel run, we then proceeded with sequencing.

**Barcode sequencing and counting**

Oxford Nanopore technology was used to perform all barcode sequencing. For stable-line experiments, where 61.2k-90k cells were sorted, we counted barcodes from three variant tests: V12-NL1a4, V30-NL2 and V37-NL1a4. From each of these tests, there was one biological sample, and three technical replicates (1199 primary PCR → 1039 nested PCR). In total, there were nine 1039-PCR pools, each of which was amplified from an independent 1199-PCR. We used the Plasmidsaurus custom sequencing service, at a 1 gigabase (Gbp) scale for each sample (9 Gbp total).

In V7-NL2a4 transient-expression experiments, each biological sample was derived from a 5k cell sort. For experiments with a standard seeding density (175k cells in 24 ww), we sequenced 3 technical replicates derived from one ratio E biological sample, and 2-4 technical replicates derived from two biological replicates of ratio F (2 tech. reps. from first sort, 4 tech. reps. from second sort). When cells were seeded at a higher cell density (HCD; 200k in 24 ww), we sequenced 3-4 technical replicates derived from two biological replicates of ratio E (4 tech. reps. from first, 3 tech. reps. from second), and 3 technical replicates derived from one biological sample of ratio F. Sequencing was performed both in-house using a MinION device, and via the Plasmidsaurus custom sequencing service at a 111 megabase scale per sample.

To count barcodes from the large sorts (61.2k-90k), we used a custom script that looked for sequences matching the designed K6x3 pattern (KKKKKK-AAGAA-KKKKKK-AGGA-KKKKKK, K=G or T) that were present in at least 2 reads in 2 or more technical replicates. Due to the large amount of data (9 Gbp total), only the following error-correction was performed inside of K6 segments: homopolymer over- and under-runs, C-T miscalls, A-G miscalls. In addition, for a barcode to be included in a technical replicate, a perfect 45 nt recombination junction sequence (5′-gagcatcccaagaggactt TC AGGTCTCCTTATTTGGGAAAACCC-3′) had to be the first, second or third most-frequently observed junction sequence of reads with the barcode. We did not count barcodes that were still non-conformant (non-K6x3) after error-correction. The barcodes and counts have been provided in Supplementary Document S1.

To compensate for lower coverage levels in 5k sort sequencing, K6x3 and non-K6x3 barcodes were counted if they were present in at least 1 read in 2 or more technical replicates, and additional error-correction was performed on barcodes before counting them. First, non-K6x3 sequences were merged with K6x3 barcodes if they had 2 or less mismatches in an alignment (1 mismatch is defined as a mismatched base, gap open, gap extension, or a base in unaligned start/end region). VSEARCH (29) was used to perform alignments with the following arguments: "--allpairs_global", "--userout", "--userfields", "query+target+alnlen+mism+gaps+opens+exts", "--acceptall". Next, remaining non-K6x3 sequences (i.e. those that were not merged into K6x3 barcodes) were clustered with other remaining non-K6x3 sequences using VSEARCH. The following arguments were used: "--cluster_size", "--consout", "--clusterout_sort", "--id 0.94", "--iddef 1", "--sizeorder", "--sizein", "--sizeout", "--uc", "—msaout". When counting barcodes, we looked for exact matches to either type of cluster sequence (K6x3 or non-K6x3) that were present in 2 or more technical replicates. The barcodes and counts have been provided in Supplementary Document S2.
